# *Haloplasma* MreB reconstitution in a minimal synthetic bacterium reveals orthogonal force directions underlying distinct Mollicute motility systems

**DOI:** 10.64898/2026.06.16.731841

**Authors:** Mone Mimura, Haruka Yuasa, Hirofumi Wada, Hana Kiyama, Yoshihito Kitaoku, Shingo Kato, Yuya Sasajima, Yuhei O Tahara, Atsuko Uenoyama, Tomoko Miyata, Fumiaki Makino, Moriya Ohkuma, Shigeyuki Kakizawa, Robert C Robinson, Keiichi Namba, Makoto Miyata

## Abstract

MreB, a bacterial actin homolog, is widely conserved in bacteria where it functions as a scaffold for a peptidoglycan synthesis. Intriguingly, members of the cell wall-lacking Mollicutes retain multiple *mreB* genes despite the absence of peptidoglycan. Previous work demonstrated that two MreB isoforms from *Spiroplasma* can reconstitute swimming motility in the minimal synthetic bacterium syn3B, indicating that MreB was repurposed for motility. Here, we expressed seven MreBs from *Haloplasma contractile*, an early-diverging Mollicute, in syn3B. Approximately 50% of cells elongated, and 15% exhibited curving and coiling movements that resembled *Haloplasma* but differed from *Spiroplasma* motility. Systematic analysis of isoform combinations revealed that movement requires specific MreB pairs, and that additional isoforms enhance motility efficiency. Cryo-electron tomography showed membrane-associated ribbons composed of MreB filaments. Mathematical modeling demonstrated that *Haloplasma*- and *Spiroplasma*-type movements can be reproduced by altering only the direction of force generation. Thus, diversification of force orientation within an actin homolog enabled independent evolution of distinct motility systems in Mollicutes.

**Teaser:** MreB a bacterial actin can drive motility of a minimal synthetic bacterium by mechanisms distinct from eukaryotic actin.

## Introduction

Actin and its superfamily, conserved in most organisms, are proteins with a molecular weight of 35-60 kDa that form filaments for which their dynamics rely on energy sources such as ATP (*1–3*). These filaments constitute a major component of the cytoskeleton and support cellular structure (*3, 4*). Moreover, in many contexts, filaments play a central role in cell motility (*5*). Actin-driven motility generally occurs through one of two well-characterized mechanisms (*5*). In the first, myosin motors slide along actin filaments, as observed in animal muscle contraction and certain forms of amoeboid motility (*6*). In the second, polymerization of actin at the leading-edge drives protrusion, as exemplified by pseudopod formation during amoeboid movement (*7*). In contrast, the bacterial actin homolog MreB does not directly generate motile force. Instead, in rod-shaped bacteria it forms membrane-associated filaments that move circumferentially in a manner coupled to peptidoglycan synthesis, functioning primarily as a scaffold for cell wall assembly (*8–10*).

However, a distinct actin-based motility mechanism has been identified in bacteria of the genus *Spiroplasma*, belonging to the class Mollicutes (*11–14*). *Spiroplasma* swim by propagating kinks along their helical cell body, effectively pushing water rearward by switching helicity from right- to left-handed and back again (*15–19*). This helix formation and switching are driven not by myosin or polymerization-based propulsion, but by multiple isoforms of a bacterial actin, MreB (*11, 20–23*). In our previous study, we demonstrated that expression of two *Spiroplasma* MreB isoforms in the minimal synthetic bacterium JCVI-syn3B (syn3B) was sufficient to induce helical cell morphology and helicity switching (*11, 24, 25*). syn3B is derived from syn3.0, a minimal synthetic bacterium constructed from *Mycoplasma mycoides*, a Mollicutes species related to *Spiroplasma* (*26, 27*). The syn3.0 genome contains only 473 genes, representing a minimal set required for autonomous proliferation. The ability of two *Spiroplasma* MreB isoforms to confer motility in this simplified cellular background suggested that *Spiroplasma* MreBs function as autonomous mechanical actuators through a mechanism distinct from canonical actin–myosin sliding or polymerization-driven protrusion (*11*).

Mollicutes, including *Spiroplasma*, evolved from Firmicutes - such as *Bacillus* and *Clostridium* - through extensive genome reduction including the loss of peptidoglycan cell wall machinery (*11–14*). In most bacteria, MreB is thought to form short filaments that spatially organize the peptidoglycan synthesis machinery, thereby maintaining rod shape (*28*). Because Mollicutes lack peptidoglycan, MreB is dispensable for its ancestral function. This raises a fundamental evolutionary question: why was MreB retained in Mollicutes, and how did it acquire novel mechanical roles absent from canonical bacterial cytoskeletons? (*11, 25, 29*).

To address this question, we focused on MreB proteins from *Haloplasma contractile* (*H*MreB), an early-diverging member of the Mollicutes (*30, 31*). *H. contractile* is a strictly anaerobic bacterium isolated from the hypersaline brine-sediment interface of the Shaban Deep in the Red Sea.

Phylogenetically, it branches at the base of the Mollicutes lineage, close to Firmicutes (*12, 30, 31*). Although its genome encodes many genes essential for peptidoglycan synthesis, no peptidoglycan layer has been detected (*30, 31*). The cells form one or two protrusions that bend and coil dynamically (*31*). The genome codes as many as seven *mreB* genes, which form phylogenetic clusters distinct from those of *Spiroplasma* (*25, 29, 30*). Here, we analyzed the evolutionary relationships of MreB proteins from *H. contractile* and its related early-diverging Mollicutes. We expressed *Haloplasma* MreB isoforms (*H*MreBs) in the minimal synthetic bacterium syn3B to dissect their functional roles. Our results demonstrate that *Haloplasma* MreBs assemble into membrane-associated ribbon structures that generate motility distinct from *Spiroplasma*, and that divergence in force directionality may underlie the evolution of alternative MreB-based motility systems within Mollicutes.

## Results

### MreBs from early-diverging Mollicutes

To investigate the evolutionary diversification of MreB within Mollicutes, we analyzed MreB sequences from early-diverging lineages and compared them with those of *Spiroplasma* and Firmicutes. Here, the term “early-diverging Mollicutes” refers to lineages that branch near the base of the Mollicutes phylogeny. MreB sequences were retrieved from the NCBI database using protein homology searches. Three MreBs were identified in *Candidatus* (*Ca*.) Xianfuyuplasma coldseepsis and *Ca.* Izimaplasma sp. HR1 and HR2. By contrast, *Haloplasma contractile* SSD-17B, *Haloplasma contractile* B5_bin.1, and Haloplasmatales bacterium (*Ca.* Inordinaticella fortuita) 4B encode seven, six, and five isoforms, respectively. These differences suggest lineage-specific expansion of MreB in *Haloplasma* species. Given this variation in copy number, we next examined the evolutionary relationships among these MreBs.

To clarify the evolutionary timing of MreB diversification, we constructed a phylogenetic tree using 33 MreB amino acid sequences derived from eight bacterial genomes, including *Bacillus subtilis* (Firmicutes) and *Spiroplasma* (Fig. 1A and fig. S1). MreBs from Mollicutes formed two distinct clades comprising early-diverging Mollicutes (clade I) and *Spiroplasma* (clade II), both clearly separated from *Bacillus subtilis*. Consistent with previous reports, *Spiroplasma* MreBs clustered into three groups (*25, 29*). Similarly, MreBs from *Haloplasma* formed three clusters, designated *H*A, *H*B, and *H*C. MreBs from *Izimaplasma* and *Xianfuyuplasma* also formed three clusters, designated *I*1, *I*2, and *I*3, although the topology within the *I*1 cluster depended on the region of amino acid sequence used. The six MreB clusters derived from early-diverging Mollicutes were further grouped into two higher-order clades: (*H*A, *I*1, *I*2) and (*H*B, *H*C, *I*3). We refer to the *Haloplasma contractile* SSD-17B seven MreB proteins as *H*MreBs B1, A1, B2, A2, C2, A3, and B4, where the letter indicates phylogenetic clade and the number indicates genome operon (Fig. 1A). Notably, although all three *Haloplasma* genomes encoded five or more MreBs, these isoforms did not form species-specific subgroups, in contrast to the clear grouping observed for *Spiroplasma* MreBs (*25, 29*). The presence of multiple conserved MreB clades across early-diverging Mollicutes suggests that MreB diversification occurred before the last common ancestor of these lineages.

**Fig. 1.**
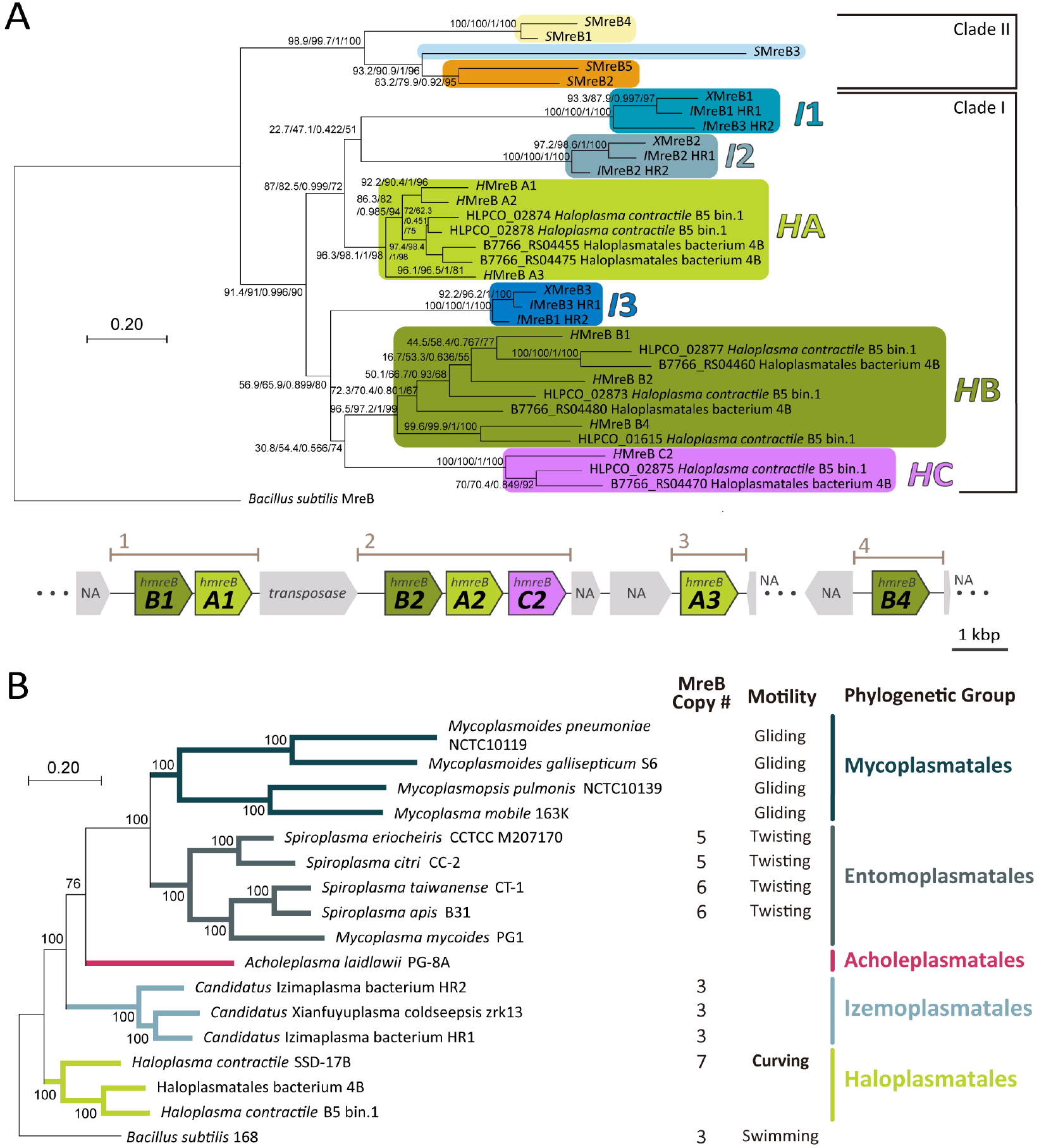
Phylogenetic relationships of Mollicutes and their MreB proteins. (**A**) Maximum-likelihood phylogenetic tree of MreB proteins from class Mollicutes. *H*MreB, *S*MreB, *X*MreB, and *I*MreB denote MreBs from *Haloplasma contractile* SSD-17B, *Spiroplasma eriocheiris*, *Candidatus* Xianfuyuplasma coldseepsis, and *Candidatus* Izimaplasma sp., respectively. Additional sequences from *Haloplasma contractile* B5 bin.1 and Haloplasmatales bacterium (*Inordinaticella fortuita*) 4B were included to resolve *Haloplasma*-related clusters. *Bacillus subtilis* MreB was used as an outgroup. Six early-diverging Mollicutes MreB clusters are color-coded and labeled (*H*A, *H*B, *H*C, *I*1, *I*2 and *I*3). Node labels on indicate support values (SH-aLRT, fast local bootstrap, approximate Bayes test, and UFBoot). Scale bar, 0.20 substitution per site. Genomic organization of *hmreB* genes in *Haloplasma contractile* SSD-17B is presented in the bottom. *hmreB B4* is coded in a distant genomic position. Genes are colored according to phylogenetic clade. (**B**) Maximum-likelihood phylogenetic tree of Mollicutes species based on concatenated alignments of 50 highly conserved coding sequences. Node labels indicate branch support assessed by UFBoot. Scale bar, 0.20 substitution per site.

To distinguish between species-level divergence and gene-specific evolution, we constructed a species phylogeny based on the 50 most conserved coding sequences from 17 species (16 Mollicutes and *Bacillus subtilis*) (Fig. 1B). *Haloplasma contractile* and related species clustered closer to Firmicutes than to *Spiroplasma*. However, MreB substitution rates relative to *Bacillus subtilis* were comparable in *Spiroplasma* (1.60 to 1.91) and *Haloplasma* (1.44 to 1.78). Thus, divergence of MreB does not strictly mirror species phylogeny, suggesting that MreB evolution may have been shaped by functional diversification rather than simply reflecting species divergence.

### *Haloplasma* MreBs cluster induces elongation and motility in syn3B

To test whether *Haloplasma* SSD-17B MreBs are sufficient to induce morphological and motility phenotypes, all seven full-length *mreB* genes were assembled into a single fragment and integrated into the *lox*P site of the syn3B genome (Fig. 2A). The resulting strain, “syn3B–Halo7”, was observed after several passages. Observation under an optical microscope revealed that approximately half of the cells were elongated, reaching up to 51 μm in length, and about 15% of the cells exhibited active movements (Fig. 2B, 4A and B, and movie S1). Hoechst staining revealed that nucleoids were distributed at intervals of 0.52 ± 0.11 µm (n = 12) throughout the entire length of the elongated cells (13.3 ± 6.8 µm, n = 12), indicating that elongation was accompanied by chromosomal reorganization along the extended axis (Fig. 2C and movie S2).

**Fig. 2.**
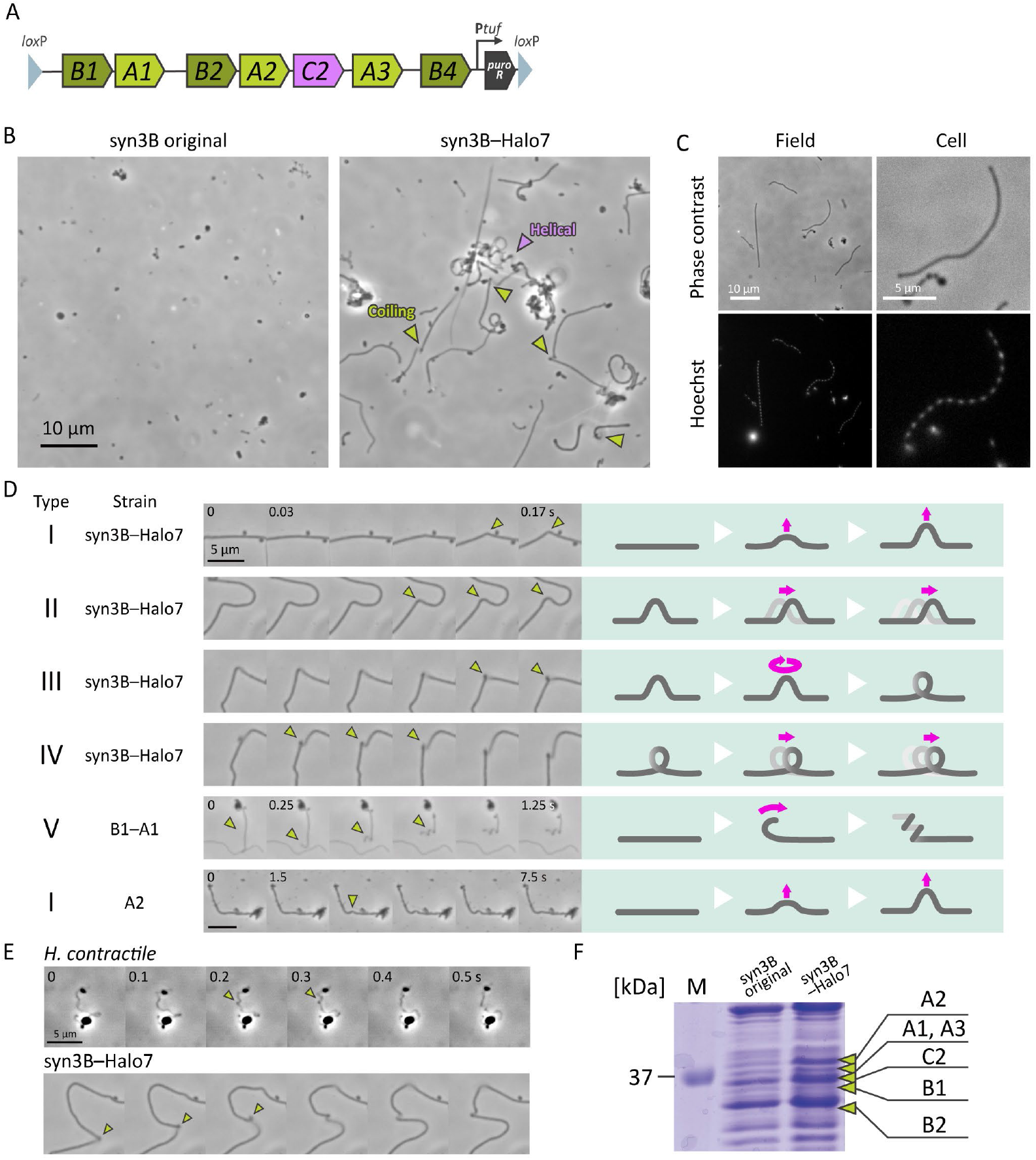
syn3B expressing seven *Haloplasma* MreBs (syn3B–Halo7). (**A**) DNA fragment used to introduce the seven *hmreB* genes into the *lox*P site in syn3B. The puromycin resistance gene (*puroR*), driven by the tuf promoter (P*tuf*), was used as a selection marker. (**B**) Phase contrast images of parental syn3B (left) and syn3B harboring the seven *hmreB* genes (syn3B–Halo7) (right) (movie S1). Coiling movements and static helices are marked by green and purple triangles, respectively. (**C**) Nucleoid positioning in elongated syn3B– Halo7 cells visualized by Hoechst staining (movie S2). (**D**) Classification of movements into five types. Consecutive video frames (0.03, 0.25 or 1.5 s intervals; left) and corresponding schematics (right) are shown. Type I–V represent curve formation, curve displacement, loop formation, loop displacement, and rolling up, respectively. Moving positions are marked by a green-colored triangle. Rare reversible type I movement was detected also in the *H*MreB A2-only strain (bottom, movie S7). *H*MreB proteins are presented only by protein numbers like A1, B2 etc. (**E**) Comparison of *Haloplasma* and syn3B–Halo7 movements by consecutive video images. Apparent displacement is marked by green-colored triangles (movie S4). (**F**) Protein profile of syn3B–Halo7 analyzed by SDS-10% PAGE stained by Coomassie staining. Lanes (left to right): Molecular weight markers, syn3B, syn3B–Halo7. Bands corresponding to *H*MreBs were identified by peptide mass fingerprinting (table S1) and marked by green arrows.

The elongated cells exhibited various movements representing a continuum of dynamic bending and translocation of curved and loop regions. The movements were classified into four patterns (Fig. 2D and 4C, and movie S3): (I) Curve formation, in which straight cells curved (observed in 100% of 87 cells); (II) Curve displacement, in which the curved region translocated (81.6%); (III) Loop formation, in which the curved region twisted to form a loop (17.2%); and (IV) Loop displacement, in which the loop translocated along the cell axis (18.4%). The movements observed in syn3B were compared with native *Haloplasma* (Fig. 2E and movie S4). *Haloplasma* exhibits tentacle-like protrusions extending from coccoid bodies (*31*). These protrusions formed transient bends and loops in some sections, qualitatively resembling the movements of syn3B– Halo7.

We next examined *Haloplasma* MreB protein expression in syn3B–Halo7 using SDS-PAGE and mass spectrometry (Fig. 2F). Distinct protein bands absent from the original syn3B cells were found in the SDS-PAGE gel within the range of 25-50 kDa. Each of the 23 bands observed in this region was analyzed by peptide mass fingerprinting (PMF) and MS/MS analysis (Fig. S2A). Six *H*MreBs were reliably detected (Mascot Score >27 and sequence coverage >25%; table S1), whilst *H*MreB B4 was not detected. These results demonstrate that expression of *Haloplasma* MreBs is sufficient to induce morphological and behavioral changes in syn3B cells.

### Functional dissection of *Haloplasma* MreB multi-isoform combinations

To investigate the contributions of individual MreB isoforms, we observed various reduced combinations of *H*MreB genes into syn3B (Fig. 3 and movies S5, S6). Strains expressing five isoforms (*H*MreBs B1–A1–B2–A2–C2 or *H*MreBs B2–A2–C2–A3–B4) exhibited cell elongation and motility similar to syn3B–Halo7 (Fig. 3A, 4A and 4B). However, the strain harboring *H*MreBs B2–A2–C2–A3–B4 exhibited significantly reduced cell length as evaluated by Student’s t-test (p<0.01, n=283 for syn3B–Halo7, n=359 for *H*MreBs B2–A2–C2–A3–B4). Expression of four *H*MreBs (B1–A1–B2–A2) also induced cell elongation and motility, although cell length was again significantly reduced relative to syn3B–Halo7 as evaluated by Student’s t-test (p<0.01, n=283 for syn3B–Halo7, n=273 for *H*MreBs B1–A1–B2–A2) (Fig. 4B). These cells exhibited an additional rolling-up pattern of movement (Type V; Fig. 2D and 4C, and movie S3).

**Fig. 3.**
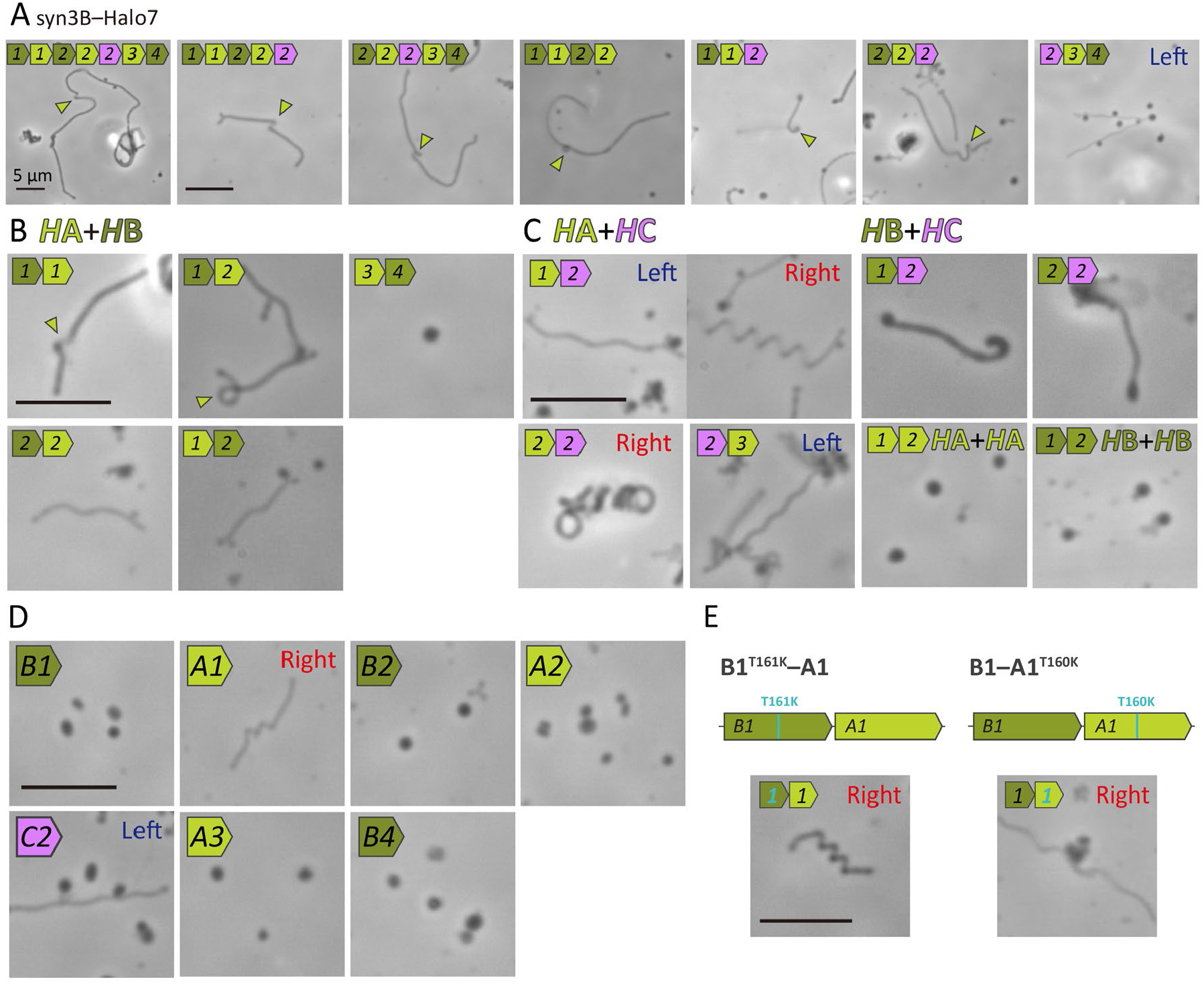
Morphology and motility of syn3B expressing defined combinations of *H*MreBs. *mreB* genes are colored light green, dark green, and purple to show MreB clusters *H*A, *H*B, and *H*C, respectively. (**A**) Cells expressing three or more *H*MreBs. Gene combinations are shown schematically in the upper-left corner of each panel. Genes are colored according to phylogenetic clades. Most combinations, except *H*MreBs B1–A1–B2–A2, include genes from all three clades (*H*A, *H*B, and *H*C) (movie S5). (**B**) Cells expressing *H*A and *H*B *H*MreB pairs. (**C**) Cells expressing *H*MreB gene pairs that did not exhibit movement. (**D**) Cells expressing a single *H*MreB (movie S6). (**E**) Cells expressing *H*MreBs B1–A1 with a mutation reducing ATPase activity.

To simplify functional comparison across clades, we generated strains containing one representative isoform from each of the *H*A, *H*B, and *H*C clades resulting in *H*MreB combinations: B1–A1–C2; B2–A2–C2; and C2–A3–B4. Elongated cells were observed in strains expressing *H*MreBs B1–A1–C2 and B2–A2–C2 (Fig. 3A, 4A and 4B). These two strains also exhibited movement patterns similar to syn3B–Halo7 (Fig. 4C), and *H*MreBs B1–A1–C2 additionally showed Type V movements (Fig. 4C). In contrast, cells expressing *H*MreBs C2–A3– B4 showed minimal elongation and no observable movement. However, a small number of cells formed a left-handed helical morphology (Fig. 3A, 4A, and 4B), potentially reflecting the absence of detectable *H*MreB B4 expression.

**Fig. 4.**
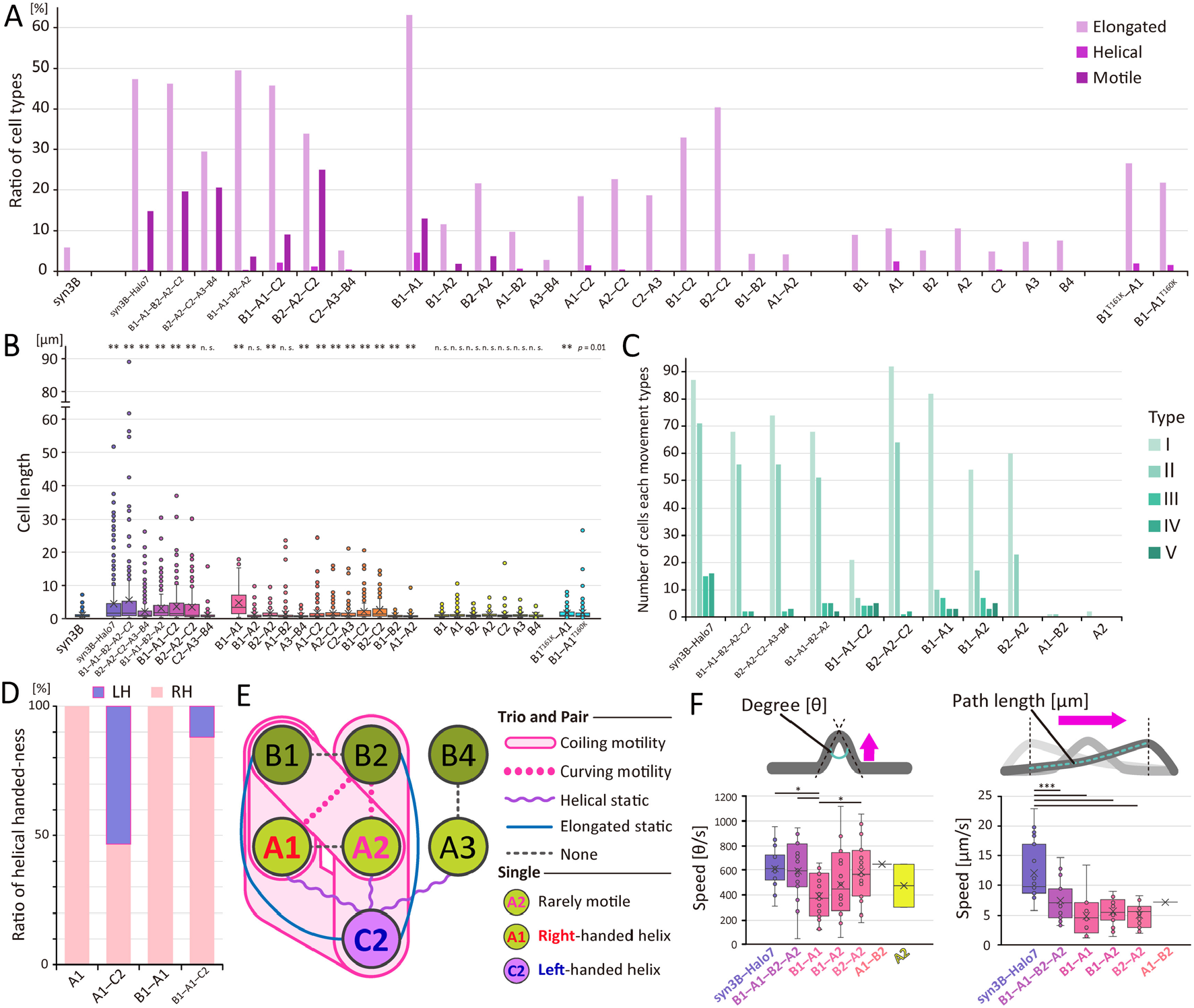
Analyses of morphology and motility of syn3B expressing *H*MreBs. *H*MreB proteins are presented only by protein numbers like A1, B2 etc. (**A**) Proportion of cells exhibiting elongation, helicity, and movement. Elongated cells include helical and motile cells. At least 118 cells were analyzed per strain. (**B**) Cell length distributions. Statistical significance was evaluated using Student’s t-test (\*\**p* < 0.01; n.s., *p* > 0.05 vs syn3B). At least 118 cells were analyzed per strain. (**C**) Number of cells exhibiting each movement type per strain. (**D**) Ratio of helical handedness. Left- and right-handedness are colored blue and red, respectively. More than 25 cells were examined for each strain. (**E**) Summary of morphological and motility features induced by different *H*MreB combinations. *H*MreBs and features are presented by circles and lines, respectively. *H*MreBs are represented by circles and phenotypic features by connecting lines. Colors indicate clades (*H*A, light green; *H*B, dark green; *H*C, purple). (**F**) Quantification of movement speeds. Speed definitions are illustrated above; measurements were performed at 1/30 s resolution, and analyzed segments are highlighted in light blue. The rate of curvature change (left) and curve displacement speed (right) are shown. Curve displacement was measured along the segment moving at constant velocity: 2.8 ± 1.1 µm (n = 19) for syn3B–Halo7; 2.1 ± 2.0 µm (n = 15) for *H*MreBs B1–A1– B2–A2; 0.57 ± 0.25 µm (n = 15) for *H*MreBs B1–A1; 0.76 ± 0.26 µm (n = 15) for *H*MreBs B1–A2; 0.51 ± 0.26 µm (n = 15) for *H*MreBs B2–A2cells, and 0.85 µm (n = 1; descriptive value only) for *H*MreBs A1–B2cells. Statistical significance was assessed using Tukey-Kramer HSD test (*** *p* < 0.002 and * *p* <0.05; n.s., *p* > 0.05), except for *H*MreBs A1–B2 and *H*MreB A2-only strains.

### Pairwise *Haloplasma* MreB combinations reveal *H*A-*H*B dependence

Given that *Spiroplasma* motility can be reduced to pairs of MreBs from different clades (*11*), we systematically tested pairwise combinations of *Haloplasma* MreBs across clades (Fig. 1A, Fig. 3B and C, and movies S5 and S6). First, we focused on combinations of *H*A and *H*B (Fig. 3B). Cells expressing *H*MreBs B1–A1 or B1–A2 exhibited significant elongation and helix formation (Fig. 3B, 4A and 4B). These strains displayed all movement types (I to V), similar to strains expressing *H*MreBs B1–A1–B2–A2 or B1–A1–C2 (Fig. 2D and 4C, and movies S3 and S5). The *H*MreBs B2–A2 strain also elongated (Fig. 3B, 4A and 4B) and exhibited curved movement but did not form loops (Fig. 4C). In contrast, *H*MreBs A1–B2 cells occasionally formed helices (Fig. 3B, 4A, and 4B), but only 1 from 4,500 cells exhibited detectable movement (Fig. 4C). The *H*MreBs A3–B4 strain showed no obvious differences from the original syn3B (Fig. 3B, 4A, and 4B), possibly reflecting the lack of detectable *H*MreB B4 expression.

We next examined *H*A and *H*C combinations (Fig. 3C). These strains elongated and occasionally formed helices but did not exhibit motility (Fig. 4A and 4B, movie S6). *H*MreBs A1–C2 strain formed both left- and right-handed helices, *H*MreBs A2–C2 cell formed right-handed helices, and the *H*MreBs C2–A3 strain formed left-handed helices, similar to *H*MreBs C2–A3–B4 (Fig. 3C). We also tested *H*B–*H*C combinations (Fig. 3C). *H*MreBs B1–C2 and B2–C2 strains elongated (Fig. 4A and 4B) but did not exhibit movement. Finally, combinations within the same clade (*H*MreBs A1–A2 and B1–B2) showed no morphological differences from the original syn3B (Fig. 3C, 4A and 4B). Together, these results indicate that motility requires specific *H*A and *H*B combinations, suggesting functional coupling between these two clades. This is mirrored in the genome structure where two pairs of *H*As and *H*Bs exist as neighboring genes, *H*MreBs B1–A1 and B2–A2 (Fig. 1A lower).

### Effects of single *Haloplasma* MreB proteins

Finally, we examined cell morphological changes and movements caused by the expression of individual *H*MreB isoforms (Fig. 3D, movies S6). *H*MreB A1-only and *H*MreB C2-only strains showed rare elongation, with 2.4% and 0.4% of cells forming right-and left-handed helices, respectively (Fig. 3D and 4A). Notably, among approximately 3,600 cells examined in the *H*MreB A2-only strain, two cells (0.056%) formed a transient curve which subsequently relaxed, a variation on movement pattern I (Fig. 2D and movie S7).

To further investigate the role of the *H*C clade protein, we introduced *H*MreB C2 into selected strains exhibiting elongation or helix formation and observed the resulting phenotypes. In *H*MreB A1-only and *H*MreBs B1–A1 cells, all helices were right-handed, however, left-handed helices appeared in *H*MreBs A1–C2 and B1–A1–C2 strains (Fig. 4D). Addition of *H*MreB C2 also resulted in helices with larger pitch and smaller width compared to *H*MreB A1-only or *H*MreBs B1–A1 strains (fig. S3). Furthermore, strains expressing *H*MreBs B2–A2 did not form loops, whereas the strain with the addition of *H*MreB C2 (B2–A2–C2) restored loop formation and increased the proportion of motile cells (Fig. 3A, 3B, 4A, and 4C). Together, these results suggest that *H*MreB C2 is not sufficient for force generation but instead modulates filament architecture, stability and efficiency. The morphological and behavioral effects of *H*MreB expression in syn3B are summarized schematically in Fig. 4E.

### ATPase activity is required for movement but not helicity

MreB proteins, including those derived from *Spiroplasma*, undergo structural changes driven by ATP hydrolysis. To determine whether ATP hydrolysis is also required for *Haloplasma* MreBs to produce motility, we mutated a conserved ATPase residue within the motile *H*MreBs B1–A1 syn3B background. We focused on a conserved threonine that is essential for ATP hydrolysis in *Spiroplasma eriocheiris* MreB5 (Thr160) (*24*). Corresponding mutations were introduced singly into each *H*MreB to produce B1^T161K^–A1 and B1–A1^T160K^ strains (Fig. 3E and fig. S4). Cells expressing either mutant elongated and formed right-handed helices but did not exhibit movement (Fig. 3E, 4A, 4B and fig. S4). Furthermore, the frequency of cell elongation and helix formation was reduced compared to the unmutated *H*MreBs B1–A1 strain (Fig. 4A). The *H*MreBs B1– A1^T160K^ strain exhibited a larger helical pitch than the *H*MreB B1–A1 strain or the *H*MreBs A1-only strain (fig. S3B). Because mutation of either *H*A or *H*B abolished motility, ATPase activity in both *H*A and *H*B subunits is required for movement, but is not for helix formation.

### Movement velocity depends on MreB composition

We measured movement speeds for three behaviors “Curve formation”, “Curve displacement”, and “Loop displacement” for representative strains (Fig. 4F and fig. S5). For curve formation, the *H*MreBs B1–A1–B2–A2 strain moved with speeds not significantly different than syn3B–Halo7 (Fig. 4F left). Among pairwise combinations, the *H*MreBs B2–A2 strain exhibited higher speeds than *H*MreBs B1–A1 strains. In contrast, curve displacement was fastest in syn3B–Halo7, which moved significantly faster than *H*MreBs B1–A1–B2–A2 or *H*MreBs B1–A1, *H*MreBs B1–A2, and *H*MreBs B2–A2 strains (Fig. 4F right). Similarly, loop displacement was faster in syn3B– Halo7 than in *H*MreBs B1–A1–B2–A2, *H*MreBs B1–A1, and *H*MreBs B1–A2 strains (fig. S5). These results indicate that reduced *H*MreB combinations generate varied speeds, but all are less efficient in sustaining coordinated displacement compared to syn3B–Halo7.

### Filamentous structures in syn3B expressing *Haloplasma* MreBs

To identify the *H*MreB structures responsible for syn3B elongation and movement, we analyzed negatively-stained samples of *H. contractile*, syn3B–Halo7 and the *H*MreBs B1–A1 strain by electron microscopy (EM) (Fig. 5A, fig. S6). The mean cell lengths of syn3B–Halo7 and *H*MreBs B1–A1 strains were 7.0 ± 2.3 μm (n = 10) and 8.8 ± 5.2 μm (n = 11), respectively, consistent with measurements obtained by optical microscopy (8.5 ± 8.8 μm, n = 134; and 6.8 ± 4.2 μm, n = 82). Cell diameters were 0.12 ± 0.06 μm (n = 10) for syn3B–Halo7 and 0.15 ± 0.05 μm (n = 15) for *H*MreBs B1–A1 strains (fig. S6, center and right). Many of *H. contractile* cells, whose normal habitat is sea water, exhibited membrane damage, likely reflecting sensitivity to low osmolarity during preparation. In partially disrupted cells, ribbon-like structures composed of filaments were exposed. Similarly, Triton-treated syn3B *H*MreB strains also revealed internal filamentous structures (Fig. 5A). Fast Fourier transform (FFT) analysis of the filament images in syn3B *H*MreB strains showed reflections corresponding to a periodicity of 5.1 nm along the filament axis and 3.4–3.5 nm in perpendicular to the filaments.

**Fig. 5.**
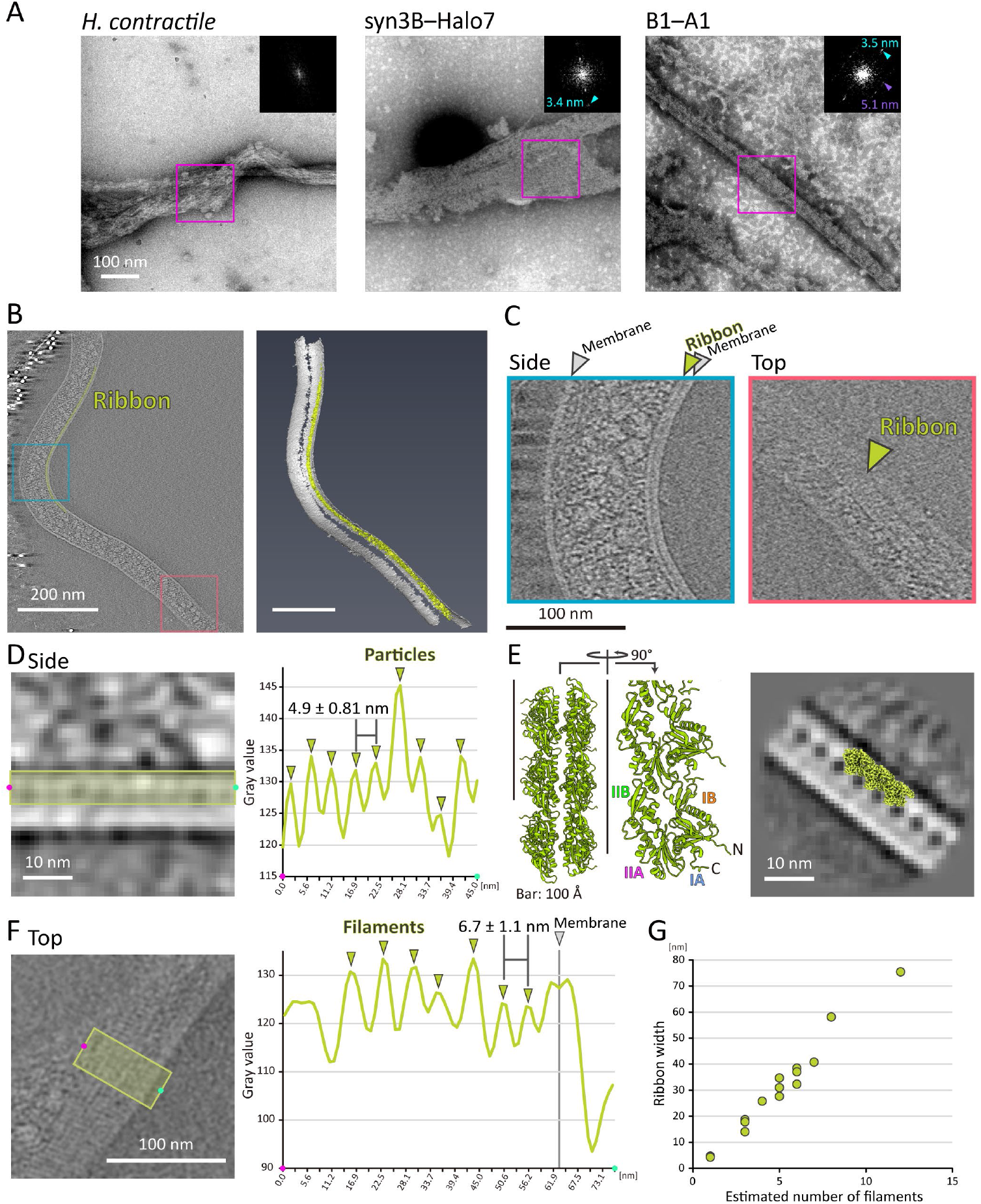
Intracellular filamentous ribbons formed by *H*MreB expression. (**A**) Filamentous structures in negatively-stained cells observed by EM. syn3B–Halo7 and *H*MreBs B1–A1 strain cells were treated by 1% Triton X-100 prior to imaging. Fast Fourier Transformation (FFT) images of boxed areas are shown (upper right). Representative diffraction spots are marked by triangles, with estimated periodicity indicated. (**B**) Cryo-electron tomogram of a *H*MreBs B1–A1 strain cell (left) and corresponding rendered image (right). Ribbon structures are highlighted in green (movie S8). (**C**) Side and top views extracted from the reconstituted volume shown in (**B**). Ribbons are marked by green triangles. (**D**) Axial periodicity of membrane-associated filaments. A smoothed tomographic slice (left) was analyzed along the green-boxed region. The mean peak-to-peak spacing was 4.9 ± 0.81 nm (data sets 47 from 6 cells). (**E**) Predicted structure of *H*MreB A1 generated by AlphaFold3 (left) and manual fitting to the averaged side-view density (right). Domain names and N- and C-termina are marked in the predicted structure. The averaged image was generated from 901 two-dimensional particles extracted from 13 tomograms of 16 cells. (**F**) Lateral periodicity of the ribbon structure. A smoothed tomographic slice (left) was profiled across the green-boxed region. The mean spacing between filament centers was 6.7 ± 1.1 nm (data sets 62 from 13 cells). (**G**) Ribbon width plotted against filament number, showing a linear relationship consistent with width increments corresponding to individual filament addition.

To further resolve filament organization, we performed cryo-electron tomography (cryo-ET) on *H*MreBs B1–A1 cells (Fig. 5B–G, movie S8). Seventeen tomograms including 23 cells were obtained. Cell thickness ranged from 63.7 to 272.7 nm, and intact membrane bilayers were observed in the section images. In well-preserved cell tomograms, a ribbon-like assembly composed of several filaments was observed immediately beneath the cell membrane on curved side of the cell (Fig. 5B and C, and movie S8). Cross-sectional views showed mainly a single layer of polymerized *H*MreB subunits on the cytoplasmic face of the membrane (Fig. 5D left), with a few regions appearing to contain two layers (fig. S7). Analysis of single layer regions revealed a periodicity of 4.9 ± 0.81 nm (data sets 47 from 6 cells) (Fig. 5D right). This value closely matches the 5.1 nm axial periodicity observed by EM of the negatively-stained cells. A two-dimensional average was generated from 901 two-dimensional particles extracted from 16 cells across 13 tomograms (fig. S8). Granular structures with dimensions similar to MreB were observed aligned along the cell membrane.

To interpret filament architecture, we predicted the structures of *H*MreB A1 and *H*MreB B1 using AlphaFold3 (*32*). *H*MreB A1 was predicted to form a filamentous assembly, although *H*MreB B1 was not (Fig. 5E left). As in canonical MreB proteins, four subdomains (IA, IB, IIA, and IIB) were identified. For fitting this model to the cryo-ET density map, residues 1–5 and 327–336 were excluded from the modeled structure, because residues 1–5, 327, and 331–336 exhibited low (70 > pLDDT > 50) or very low confidence (pLDDT < 50). The fitting suggested that the IA–IB domain faces the membrane (Fig. 5E right; fig. S9), guided by the presence of a protrusion on the IA–IB domain which is absent from the IIA–IIB domain.

Reconstitution of the ribbon structure from the top view revealed multiple aligned filaments (Fig. 5F). The spacing between adjacent filaments was 6.7 ± 1.1 nm (data sets 62 from 13 cells) (Fig. 5F), approximately twice the transverse periodicity observed with negatively-stained samples by EM. This discrepancy likely reflects differences in contrast: individual filaments are resolved under high-contrast negative staining, whereas cryo-ET primarily visualizes double-stranded assemblies. The number of filaments within a ribbon ranged from 1 to 12, and ribbon width increased proportionally with filament number, producing a linear relationship (17 data sets from 16 cells) (Fig. 5G). This indicates that the ribbon consists of aligned filaments arranged side-by-side with a constant inter-filament spacing. These structural observations provide a physical basis for *H*A-*H*B-dependent helicity and motility in syn3B.

### Difference between *Haloplasma* and *Spiroplasma* motilities

In *H*MreB-mediated syn3B movements, initially straight cells formed localized curves that subsequently migrated and expanded laterally, giving rise to loops and coils. This mode of movement differs fundamentally from that induced by *Spiroplasma* MreBs, in which the handedness of a pre-existing helix is inverted during propagation (Fig. 6A). Closer analysis suggested that *H*MreB-driven motility involves axial force generation, while *Spiroplasma* MreB-driven motility is based on shear force. To test this hypothesis, we performed mathematical modeling of the movement types (Fig. 6B and movie S9; see Supplementary Text) (*17, 18*). The model assumes that (i) changes in axial curvature are driven by axial force, whereas changes in twist (perpendicular to the axis) are driven by shear force; (ii) these states transition in a sigmoidal manner; and (iii) state transitions propagate cooperatively along the cell axis. Using these assumptions, the model successfully reproduced the *Spiroplasma* motility, as previously demonstrated (*17, 18*). Remarkably, by keeping all parameters constant and changing only the direction of force generation, the model reproduced *H*MreB-driven motility. These results indicate that the distinct motility modes of *Haloplasma* and *Spiroplasma* arise primarily from differences in force directionality. Thus, directional control of force generation appears to be a key evolutionary variable underlying diversification of MreB-based motility systems.

**Fig. 6.**
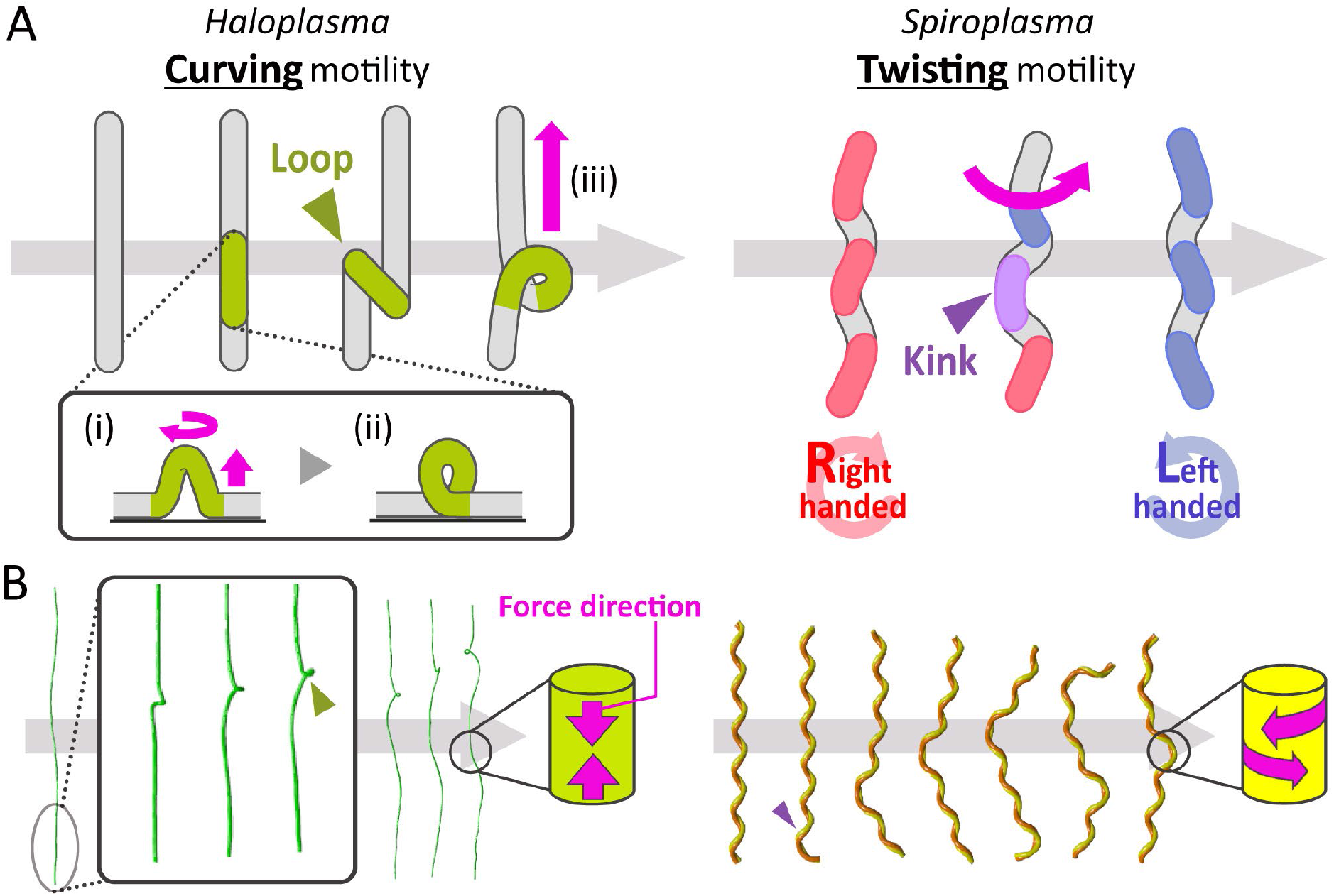
Distinct force orientations underlie *Haloplasma*- and *Spiroplasma*-type motilities reconstructed in syn3B. (**A**) Comparison of movement modes generated by *Haloplasma* and *Spiroplasma* MreBs in syn3B. Movement directions are indicated by magenta arrows. *Haloplasma*-type motility proceeds in three steps: (i) local curvature develops in an initially linear segment with minor twisting; (ii) continued twisting generates a loop; (iii) the loop travels along the cell axis. In contrast, *Spiroplasma*-type motility involves helicity switching between right- and left-handed states, accompanied with the kink traveling along the helical body (movie S9). (**B**) Mathematical modeling of the two motility types. The model incorporates three assumptions: (i) spontaneous sigmoidal transitions between two structural states; (ii) cooperative propagation of state transitions along the cell axis; and (iii) Change in curvature driven by axial force (*Haloplasma*) or twist driven by shear force (*Spiroplasma*). Altering only the direction of force generation was sufficient to reproduce both motility types.

## Discussion

Expression of *H*MreB genes in syn3B elongated the cells and enabled lateral bending movements relative to the cell axis, reminiscent of *Haloplasma* cell behaviors, demonstrating that *H*MreBs are a complete motility system. The cells contained a ribbon-like structure extending to both cell poles beneath the membrane. These results indicate that *H*MreBs confer a distinct type of motility from that of *Spiroplasma*, mediated through the formation of membrane associated filaments.

### Roles of *Haloplasma* MreB proteins

The MreB proteins of *Haloplasma* species were classified phylogenetically into three groups: *H*A, *H*B, and *H*C (Fig. 1A). Combinations of *H*A and *H*B consistently induced motility in syn3B (Fig. 3B and 4A), where other pairings did not. This result resembles the case of *Spiroplasma*, where motility occurs through specific combinations of *S*MreBs (*11*). syn3B cells expressing *H*A or *H*B alone occasionally exhibited elongation (Fig. 3D and 4A), suggesting partial MreB functional autonomy. The cells expressing *H*MreB A1 formed coil, whereas those with *H*MreB B1 did not. These observations raise the possibility that *H*A-type MreBs form relatively stable coils, whereas *H*B-type MreBs contribute primarily to force generation along the axis. This interpretation parallels findings in *Spiroplasma*, where MreB5 forms relatively stable filaments *in vitro* (*21, 24, 33, 34*).

Although various movements occurred with combinations of two *H*MreB types, strains expressing seven *H*MreBs generally exhibited higher velocities and more sustained displacement than those expressing only pairs (Fig. 4F). This suggests that while *H*A and *H*B represent the minimal functional unit for motility, additional *H*MreB isoforms enhance mechanical efficiency, coordination, or structural stability (*11*). Cryo-ET results indicated that elongation and curvature of syn3B and *Haloplasma* cells are associated with a ribbon composed of several *H*MreB filaments arranged in aligned beneath the membrane. This organization resembles the ribbon formed by several *S*MreBs in *Spiroplasma* that generates the swimming motility. However, the relative orientation of *H*A- and *H*B-type filaments could not be resolved due to limited resolution. The ribbon appeared mainly as a single layer beneath the membrane, although it remains possible that additional layers form transiently during force generation. This is because if force arises between the two layers of *H*MreB filaments, it is expected to cause the bending observed in this study. If bending arises from differential strain between adjacent filaments or layers, such multilayer interactions could account for curvature generation. Further structural analysis will be required to test this hypothesis.

### Differences and evolutionary development of *Haloplasma* and *Spiroplasma* motilities

Mathematical modeling demonstrated that the difference between *Haloplasma*- and *Spiroplasma*-type can be explained primarily by the direction of force generation (Fig. 6, movie S9). Remarkably, changing only force direction while keeping all other parameters constant reproduced the two distinct motility modes.

Phylogenetic analysis indicates that the MreB proteins of *Haloplasma* and *Spiroplasma* evolved independently from those of Firmicutes, represented by *Bacillus subtilis* (Fig. 1A). Thus, the immediate ancestors of these two motility systems were unlikely to share a motile common ancestor. Taken together, these results suggest that acquisition of force-generating capacity from a canonical MreB scaffold may be relatively accessible evolutionarily. This interpretation is consistent with the rare but detectable movement observed in the syn3B strain expressing *H*MreB A2 alone (Fig. 2D, 4C; movie S7). Such residual activity may reflect retention of latent mechanical properties in ancestral MreB proteins prior to full specialization. In this scenario, modest mutations altering filament geometry or ATPase-coupled conformational changes could convert a static Firmicutes MreB scaffold into an actuator.

Several members of actin superfamily are involved in transport and movement, typically through harnessing polymerization and depolymerization dynamics (*35, 36*). In contrast, the behaviors of *Haloplasma* and *Spiroplasma* MreBs in syn3B do not appear to directly reflect simple polymerization-driven processes. For MreBs, movement likely involves an element of coordinated conformational changes within stable filament assemblies (*11, 33, 37, 38*). Distinguishing between polymerization, cooperative conformational switching, and inter-filament strain generation contributions to movement will require future biophysical experiments.

### syn3B as a minimal cellular platform

In this and previous studies (*11*), syn3B was used to investigate how MreB proteins induces changes in cell morphology and behavior. Beyond this specific application, syn3B represents a useful platform for studying protein-driven cellular mechanics in a simplified background. The elongated morphology induced by *H*MreB expression is particularly advantageous. Strains expressing five *H*MreBs B1–A1–B2–A2–C2 or the full cluster (syn3B–Halo7) reached lengths of up to ∼50 µm (Fig. 2B, 3A and 4B). Elongation likely occurs because the submembrane MreB ribbon interferes with normal cell division. Several elongated strains also exhibited motility, which may complicate protein tracking experiments. However, strains expressing *H*MreBs B1– C2 or B2–C2 remained relatively stable, with 33% and 40% of cells elongating to maximum lengths of 20.6 or 15.1 μm, respectively. These strains may therefore provide practical tools for imaging and intracellular organization studies.

Interestingly, Hoechst staining revealed regularly spaced nucleoids along elongated syn3B–Halo7 cells (Fig. 2C). In many bacteria, chromosomal positioning is mediated by the Par system (*39*). Because syn3B lacks canonical partitioning systems, this observation suggests the presence of an alternative, possibly mechanical or geometry-dependent mechanism for DNA distribution. The MreB ribbon may contribute indirectly to genome positioning by inhibiting cell division at a specific stage of DNA segregation. Thus, these cells may be useful for studying novel aspects of DNA segregation.

### Origin of motility

Motility is a defining feature of living organisms and plays a central role in survival (*36*). Cell motilities are classified into 18 mechanisms based on factors such as the structure of force-generation units. Many motility mechanisms are structurally complex and difficult to trace to immediate ancestors, likely reflecting rapid evolutionary diversification. In contrast, Mollicutes provide rare examples in which motility systems can be linked to pre-existing cellular machinery.

For example, the gliding motor of *Mycoplasma mobile* (“G_1_-type rotary ATPase”) is derived from F_1_-type rotary ATPase (*40–42*), and *Spiroplasma* swimming motility originated from the MreB scaffold associated with peptidoglycan synthase (*28*). The present study adds a further example: *Haloplasma*, one of the earliest-diverging Mollicutes (*12, 30, 31*), evolved an MreB actuator distinct from that of *Spiroplasma*. Based on these observations, we propose that motility can emerge when intrinsic mechanical fluctuations of essential proteins become amplified, coordinated, and transmitted to the cell surface (*36*). In this view, motility systems need not arise de novo but can evolve from subtle mechanical properties already embedded within housekeeping proteins. The diversification of *Haloplasma* and *Spiroplasma* MreBs illustrates how modification of force directionality alone may be sufficient to generate fundamentally distinct movement strategies. Thus, MreB provides a compelling example of how a protein originally dedicated to cell maintenance can be repurposed into a motility actuator through evolutionary refinement.

## Materials and Methods

### Phylogenetic analysis

The MreB phylogenetic tree was constructed using sequences identified as MreB homologs in NCBI database (https://www.ncbi.nlm.nih.gov/). MreB sequences of *Candidatus* Xianfuyuplasma coldseeps were extracted from the annotated genome found in NCBI. MreB sequences of *H. contractile* B5_bin.1 were not found in NCBI. Therefore, a BLAST search (https://blast.ncbi.nlm.nih.gov/Blast.cgi) was performed using seven *H*MreB sequences as queries from strain SSD-17B, and the annotated scaffold sequences as subjects. Due to the lack of genome annotation for *H. contractile* B5_bin.1 in NCBI genome database, coding sequences (CDSs) were annotated using Prokka (*43*) with the default settings. Sequences were extracted as a fasta-formatted file using a Python 3.11.1 script with the aid of the Biopython library (*44*). MreB sequences covered query more than 98% were used for the tree construction.

Genome sequences of Mollicutes were downloaded from NCBI genome database (https://www.ncbi.nlm.nih.gov/home/genomes/). Phylogenetic analysis was conducted based on 50 CDSs with the highest conservation among class Mollicutes, selected according to their sequence identities to homologs in *H. contractile* SSD-17B. The selected CDSs are listed in the supporting data (Data set S1). Multiple sequence alignments were generated using MUSCLE (*45*). Alignments were trimmed using trimAl with the automated1 option (*46*). Trimmed alignments were concatenated using AMAS with the concat command (*47*). Phylogenetic trees were constructed using the maximum-likelihood method implemented in IQ-TREE using the concatenated sequences (*48*). For the MreB tree, the Q yeast matrix with discrete Gamma model (4 rate categories) best fit the data according to AIC, AICc, and BIC criteria. For the concatenated protein tree, the LG substitution matrix with a proportion of invariable sites and four FreeRate categories (*49, 50*) was selected. Node support values were estimated using the SH-like approximate likelihood ratio test (*51*), fast local bootstrap probability method (*52*), approximate Bayes test (*53*), and ultrafast bootstrap approximation (*54, 55*).

### Bacterial strains

JCVI-syn3B (GenBank, CP069345.1) (*26, 27*), *Haloplasma contractile* SSD-17B strain (GenBank, GCA_000215935.3) (*30, 31*) and *Escherichia coli* (DH5α and EPI400) were cultured in SP4, A-MMX and LB media at 37, 30, and 37°C, respectively, with appropriate antibiotics if necessary.

### Constructing syn3B strains

DNA fragments *hmreB B1–A1* and *hmreB A3* were chemically synthesized (GenScript, Piscataway, USA). Other *HmreB* DNA fragments were amplified by PCR using *Haloplasma* cells. *puroR* (puromycin resistance gene) and vector fragments were amplified from pSD079 by PCR. Primer sequences are listed as table S2. DNA fragments were assembled using the In-Fusion HD Cloning Kit (Takara Bio Inc., Kusatsu, Japan). The assembled products were amplified in *E. coli* DH5α for pHcNb1a1, and EPI400 for pHcNa3 (fig. S10). EPI400 was used to solve poor yield problems. Other DNA fragments were generated by an *in vitro* assembly (*56*), using NEBuilder HiFi assembly® (fig. S10). syn3B transformation was performed as previously described (*11, 57*).

### Protein profile and identification

Cellular proteins were separated by SDS-PAGE and identified by Mascot search version 2.5.1, as previously described (*11, 58*), using databases JCVI-syn3B (taxonomy ID: 2806337) and *H. contractile* (taxonomy ID: 1033810).

### Optical microscopy

syn3B cells were collected and suspended in PBS (68.4 mM NaCl, 57.9 mM Na_2_HPO_4_, 16.9 mM NaH_2_PO_4_·2H_2_O) including 0.5% methylcellulose and observed in a tunnel slide, as previously described (*11, 59*). For genome visualization, the cultured cells were stained by 0.56 mM Hoechst 33342 (Nacalai Tesque, Kyoto, Japan) at 37°C for 30 min. After insertion into a tunnel slide, the solution was replaced by PBS containing 0.1% methylcellulose. *H. contractile* cells were transferred under anaerobic conditions (1 µL cell cultures) (*31*), placed between coverslips under the air conditions, and sealed by nail polish. Cells were observed using an IX71 inverted microscope (Evident, Tokyo, Japan) equipped with a UPlanSApo 100× 1.4 numerical aperture Ph3 and complementary metal-oxide semiconductor (CMOS) camera, DMK33UX174 (The Imaging Source Asia Co. Ltd. Taipei, Taiwan). Videos were analyzed by ImageJ ver. 1.53k (*60*) and 2.14.0/1.54f (Fiji) (*61*) with the ObjectJ plugin (https://sils.fnwi.uva.nl/bcb/objectj/index.html) for morphological changes.

### Electron microscopy

Negative-staining method and Triton treatment were performed as previously described (*11*). For cryo-EM, 3.0 μl of sample solution containing 10 nm colloidal gold was applied onto a glow-discharged holey carbon grid (Quantifoil R1.2/1.3 Cu grid, Quantifoil Micro Tools, Germany). Grids were plunge-frozen into liquid ethane using a Vitrobot mark IV (Thermo Fisher Science, Waltham, MA, USA) with 3 s blotting at 4°C and 90% humidity. Data was collected using a CRYO ARM 300 II (JEOL, Akishima, Japan) equipped with a cold field-emission electron gun operated at 300 kV, an Ω-type energy filter with a 20 eV slit width, and a K3 Summit direct electron detector camera (Gatan, Pleasanton, USA). Movie frames were recorded at ×25,000 magnification (pixel size of 2.343 Å) with a target defocus from −5.0 μm. Tilt series were acquired from -60° to 60° with a dose-symmetric tilt scheme at 3° increments using serialEM 4.0.

### Tomogram image processing

Motion correction was performed in RELION 4.0 (*62*). Tilt alignment and 4× binned tomogram generation were carried out using IMOD 4.11 (*63*). Reconstruction used the SIRT algorithm. Contrast adjustment of the tomograms was performed in Fiji. Tomograms were manually segmented using Amira 2019.4 (Thermo Fisher Scientific) using the threshold tool. A supplementary movie was generated using Amira. To measure the side and top widths of the ribbon, the images were processed twice with the Smooth function in Fiji. Line widths were set to 6 pixels (5.6232 nm) and 45 pixels (42.174 nm), respectively, and measurements were obtained using the Line Plot function. For cell diameter measurements, a line width of 20 pixels (18.744 nm) was used.

### 2D averaging and manual fitting

Cross-sectional views were extracted from the reconstructed tomograms, and particles were manually picked using RELION 4.0. 2D classification was performed three times to obtain optimal class averages. *H*MreBs A1 and B1 predicted structures containing six subunits were generated individually using the AlphaFold3 server (https://alphafoldserver.com/) (*32*). The structure of *H*MreB A1 predicted as antiparallel double filaments was output using ChimeraX (*64*). Protofilaments were manually fitted into 2D averaged cryo-EM-derived images.

### Mathematical modeling

Cell shape dynamics were modeled by representing the bacterium as an inextensible elastic rod with bending and twisting energies governed by Hookean elasticity. *Haloplasma*- and *Spiroplasma*-type motilities were reproduced by imposing a propagating preferred curvature or twist, respectively, and numerically integrating the overdamped Langevin equations of motion. Specific details can be found in the Supplementary text.

## Supporting information

Supplemental file and will be used for the link to the file on the preprint.

## Acknowledgments

We thank A. Antunes at University of Saint Joseph, Macau for helpful discussion and T. Shimonaka at the Osaka Metropolitan University for the technical assistance with mass spectrometry.

## Author contributions

Conceptualization: Mimura, Miyata

Phylogenetic tree: Mimura, YK

Methodology: Mimura, HK, AU, Kato, Kakizawa

Investigation: Mimura

Conventional electron microscopy: YOT

Cryo-electron microscopy: HY, YS, TM, FM, KN

Mathematical modelling: HW

Supervision: Kakizawa, RR, KN, Miyata

Writing—original draft: Mimura, HY, Miyata

Writing—review & editing: Mimura, HY, HK, YK, YOT, SK, RR, KN, Miyata

## Data and materials availability

The cryo-EM map is available on the Electron Microscopy Data Bank (EMDB) under accession code: EMD-83100. All other data needed to evaluate the conclusions in the paper are present in the paper and/or the Supplementary Materials.

## References

1. R. Dominguez, K. C. Holmes, Actin structure and function. Annu Rev Biophys 40, 169–186 (2011).

2. H. M. Meir et al., Neuroblastoma. Saudi Med J 22, 674–680 (2001).

3. T. D. Pollard, J. A. Cooper, Actin, a central player in cell shape and movement. Science 326, 1208–1212 (2009).

4. D. A. Fletcher, R. D. Mullins, Cell mechanics and the cytoskeleton. Nature 463, 485–492 (2010).

5. T. D. Pollard, G. G. Borisy, Cellular motility driven by assembly and disassembly of actin filaments. Cell 112, 453–465 (2003).

6. M. A. Hartman, J. A. Spudich, The myosin superfamily at a glance. J Cell Sci 125, 1627–1632 (2012).

7. A. Mogilner, G. Oster, Cell motility driven by actin polymerization. Biophys J 71, 3030–3045 (1996).

8. J. Dominguez-Escobar et al., Processive movement of MreB-associated cell wall biosynthetic complexes in bacteria. Science 333, 225–228 (2011).

9. E. C. Garner et al., Coupled, circumferential motions of the cell wall synthesis machinery and MreB filaments in B. subtilis. Science 333, 222–225 (2011).

10. S. van Teeffelen et al., The bacterial actin MreB rotates, and rotation depends on cell-wall assembly. Proc Natl Acad Sci U S A 108, 15822–15827 (2011).

11. H. Kiyama, S. Kakizawa, Y. Sasajima, Y. O. Tahara, M. Miyata, Reconstitution of a minimal motility system based on *Spiroplasma* swimming by two bacterial actins in a synthetic minimal bacterium. Sci Adv 8, eabo7490 (2022).

12. Y. Wang et al., Phylogenomics of expanding uncultured environmental Tenericutes provides insights into their pathogenicity and evolutionary relationship with *Bacilli*. BMC Genomics 21, 408 (2020).

13. H. Grosjean et al., Predicting the minimal translation apparatus: lessons from the reductive evolution of mollicutes. PLoS Genet 10, e1004363 (2014).

14. S. Razin, D. Yogev, Y. Naot, Molecular biology and pathogenicity of mycoplasmas. Microbiol Mol Biol Rev 62, 1094–1156 (1998).

15. Y. Sasajima, M. Miyata, Prospects for the mechanism of *Spiroplasma* swimming. Front Microbiol 12, 706426 (2021).

16. D. Nakane, T. Ito, T. Nishizaka, Coexistence of two chiral helices produces kink translation in *Spiroplasma* swimming. J Bacteriol 202, e00735–00719 (2020).

17. H. Wada, R. R. Netz, Hydrodynamics of helical-shaped bacterial motility. Phys Rev E Stat Nonlin Soft Matter Phys 80, 021921 (2009).

18. H. Wada, R. R. Netz, Model for self-propulsive helical filaments: kink-pair propagation. Phys Rev Lett 99, 108102 (2007).

19. J. W. Shaevitz, J. Y. Lee, D. A. Fletcher, *Spiroplasma* swim by a processive change in body helicity. Cell 122, 941–945 (2005).

20. C. Lartigue et al., Cytoskeletal components can turn wall-less spherical bacteria into kinking helices. Nat Commun 13, 6930 (2022).

21. S. Harne et al., MreB5 Is a determinant of rod-to-helical transition in the cell-wall-less bacterium *Spiroplasma*. Curr Biol 30, 4753–4762.e4757 (2020).

22. S. Trachtenberg et al., Structure of the cytoskeleton of *Spiroplasma melliferum* BC3 and its interactions with the cell membrane. J Mol Biol 378, 778–789 (2008).

23. J. Kürner, A. S. Frangakis, W. Baumeister, Cryo-electron tomography reveals the cytoskeletal structure of *Spiroplasma melliferum*. Science 307, 436–438 (2005).

24. D. Takahashi et al., Structure and polymerization dynamics of bacterial actin MreB3 and MreB5 involved in *Spiroplasma* swimming. Open Biol, in press (2022).

25. C. Ku, W. S. Lo, C. H. Kuo, Molecular evolution of the actin-like MreB protein gene family in wall-less bacteria. Biochem Biophys Res Commun 446, 927–932 (2014).

26. J. F. Pelletier et al., Genetic requirements for cell division in a genomically minimal cell. Cell 184, 2430–2440 e2416 (2021).

27. C. A. Hutchison, 3rd et al., Design and synthesis of a minimal bacterial genome. Science 351, aad6253 (2016).

28. H. Shi, B. P. Bratton, Z. Gitai, K. C. Huang, How to build a bacterial cell: MreB as the foreman of *E. coli* construction. Cell 172, 1294–1305 (2018).

29. D. Takahashi, I. Fujiwara, M. Miyata, Phylogenetic origin and sequence features of MreB from the wall-less swimming bacteria *Spiroplasma*. Biochem Biophys Res Commun 533, 638–644 (2020).

30. A. Antunes et al., Genome sequence of *Haloplasma* contractile, an unusual contractile bacterium from a deep-sea anoxic brine lake. J Bacteriol 193, 4551–4552 (2011).

31. A. Antunes et al., A new lineage of halophilic, wall-less, contractile bacteria from a brine-filled deep of the Red Sea. J Bacteriol 190, 3580–3587 (2008).

32. J. Abramson et al., Accurate structure prediction of biomolecular interactions with AlphaFold 3. Nature 630, 493–500 (2024).

33. D. Takahashi, H. Kiyama, H. T. Matsubayashi, I. Fujiwara, M. Miyata, A bacterial actin with high ATPase activity regulates the polymerization of a partner MreB isoform essential for *Spiroplasma* swimming motility. J Biol Chem 301, 110462 (2025).

34. D. Takahashi, M. Miyata, I. Fujiwara, Assembly properties of bacterial actin MreB involved in *Spiroplasma* swimming motility. J Biol Chem 299, 104793 (2023).

35. L. K. Fritz-Laylin, M. A. Titus, The evolution and diversity of actin-dependent cell migration. Mol Biol Cell 34, 7 (2023).

36. M. Miyata et al., Tree of motility - A proposed history of motility systems in the tree of life. Genes Cells 25, 6–21 (2020).

37. T. Tanaka, H. Kiyama, Y. V. Morimoto, T. Nishizaka, M. Miyata, Live imaging of bacterial actin MreBs from *Spiroplasma* causing helicity switching of a minimal synthetic cell. Biophys Physicobiol 23, e230017 (2026).

38. D. Takahashi, M. Miyata, I. Fujiwara, Cooperative and divergent properties of bacterial actin isoforms in *Spiroplasma* swimming. Biophys Physicobiol 23, e230012 (2026).

39. X. Wang, P. Montero Llopis, D. Z. Rudner, Organization and segregation of bacterial chromosomes. Nat Rev Genet 14, 191–203 (2013).

40. T. Toyonaga et al., Dimeric assembly of F(1)-like ATPase for the gliding motility of *Mycoplasma*. Sci Adv 11, eadr9319 (2025).

41. T. Toyonaga et al., Chained structure of dimeric F_1_-like ATPase in *Mycoplasma mobile* gliding machinery. mBio 12, e0141421 (2021).

42. M. Nishikawa et al., Refined mechanism of *Mycoplasma mobile* gliding based on structure, ATPase activity, and sialic acid binding of machinery. mBio 10, e02846–02819 (2019).

43. T. Seemann, Prokka: rapid prokaryotic genome annotation. Bioinformatics 30, 2068–2069 (2014).

44. P. J. Cock et al., Biopython: freely available Python tools for computational molecular biology and bioinformatics. Bioinformatics 25, 1422–1423 (2009).

45. R. C. Edgar, MUSCLE: multiple sequence alignment with high accuracy and high throughput. Nucleic Acids Res 32, 1792–1797 (2004).

46. S. Capella-Gutierrez, J. M. Silla-Martinez, T. Gabaldon, trimAl: a tool for automated alignment trimming in large-scale phylogenetic analyses. Bioinformatics 25, 1972–1973 (2009).

47. M. L. Borowiec, AMAS: a fast tool for alignment manipulation and computing of summary statistics. PeerJ 4, e1660 (2016).

48. B. Q. Minh et al., IQ-TREE 2: New Models and Efficient Methods for Phylogenetic Inference in the Genomic Era. Mol Biol Evol 37, 1530–1534 (2020).

49. J. Soubrier et al., The influence of rate heterogeneity among sites on the time dependence of molecular rates. Mol Biol Evol 29, 3345–3358 (2012).

50. S. Q. Le, O. Gascuel, An improved general amino acid replacement matrix. Mol Biol Evol 25, 1307–1320 (2008).

51. S. Guindon et al., New algorithms and methods to estimate maximum-likelihood phylogenies: assessing the performance of PhyML 3.0. Syst Biol 59, 307–321 (2010).

52. J. Adachi, MOLPHY version 2.3 : programs for molecular phylogenetics based on maximum likelihood. Comp Sci Monogr 28, 1–150 (1996).

53. M. Anisimova, M. Gil, J. F. Dufayard, C. Dessimoz, O. Gascuel, Survey of branch support methods demonstrates accuracy, power, and robustness of fast likelihood-based approximation schemes. Syst Biol 60, 685–699 (2011).

54. D. T. Hoang, O. Chernomor, A. von Haeseler, B. Q. Minh, L. S. Vinh, UFBoot2: Improving the Ultrafast Bootstrap Approximation. Mol Biol Evol 35, 518–522 (2018).

55. B. Q. Minh, M. A. Nguyen, A. von Haeseler, Ultrafast approximation for phylogenetic bootstrap. Mol Biol Evol 30, 1188–1195 (2013).

56. A. Uenoyama, H. Kiyama, M. Mimura, M. Miyata, Rapid in vitro method to assemble and transfer DNA fragments into the JCVI-syn3B minimal synthetic bacterial genome through Cre/loxP system. Biophys Physicobiol 21, e210024 (2024).

57. B. J. Karas et al., Rescue of mutant fitness defects using in vitro reconstituted designer transposons in Mycoplasma mycoides. Front Microbiol 5, 369 (2014).

58. M. Yabe et al., Assembly formation of P65 protein, featured by an intrinsically disordered region involved in gliding machinery of *Mycoplasma pneumoniae*. Biomolecules 15, (2025).

59. P. Liu et al., Chemotaxis without conventional two-component system, based on cell polarity and aerobic conditions in helicity-switching swimming of *Spiroplasma eriocheiris*. Front Microbiol 8, 58 (2017).

60. C. A. Schneider, W. S. Rasband, K. W. Eliceiri, NIH Image to ImageJ: 25 years of image analysis. Nat Methods 9, 671–675 (2012).

61. J. Schindelin et al., Fiji: an open-source platform for biological-image analysis. Nat Methods 9, 676-682 (2012).

62. S. H. Scheres, RELION: implementation of a Bayesian approach to cryo-EM structure determination. J Struct Biol 180, 519–530 (2012).

63. J. R. Kremer, D. N. Mastronarde, J. R. McIntosh, Computer visualization of three-dimensional image data using IMOD. J Struct Biol 116, 71–76 (1996).

64. E. F. Pettersen et al., UCSF ChimeraX: Structure visualization for researchers, educators, and developers. Protein Sci 30, 70–82 (2021).

