## Supplemental file and will be used for the link to the file on the preprint. for "*Haloplasma* MreB reconstitution in a minimal synthetic bacterium reveals orthogonal force directions underlying distinct Mollicute motility systems": Supp Halosyn3B_202600805-2.pdf

Mone Mimura *et al.*

**This PDF file includes:**

Supplementary Text
Figs. S1 to S11
Tables S1 to S3
Legends for movies S1 to S9
Legends for data set S1
References (65 to 69)

**Other Supplementary Materials for this manuscript include the following:**
[use this section only if you have movies, audio or data files]
Movies S1 to S9
Data set S1

Color Align Properties results

```
G, A, V, L, I
F, Y, W
C, M
S, T
K, R, H
D, E
N, Q
P
```

```
SMreB1_DSM21848        ----------MALINNKKPTFVSIDLGTAFTLVYISGSGIVYNEPSIVAYKIKENRIIAVGAEAYKMIGKGNKSIRIVRP 70
SMreB2_DSM21848        ---------MANYKFGKEYSFLALDLGTANTVAYVAGQGIVYNEPSMMAYDTLSNSLVALGEEAYKMIGKTHDHIKMVTP 71
SMreB3_DSM21848        MTITDVLKNTFNISPKPPRKFIAIDLGTTNSIAYIGGRGIIYNEASVMAYETGTKKLVALGEDARKLIGKTHDKIEIYTP 80
SMreB4_DSM21848        -------MAGFNSGKSKRPTFVSMDLGTANTLVYVSGSGVVYNEPSIVAYRIKENRIIAVGIEAYKMIGKGNKSIRIVRP 73
SMreB5_DSM21848        --------------MKPERPFISLDLGTANVLAYVSGQGIIYNEPSLMAYDTKTNKLIALGKEAYDMIGKTHDQIRMVTP 66
XMreB1_zrk13           -----------MSEKPKGSFNVGVDLGTSNLLIYVEGRGTIFNEPSYIAVDKATKQVVSVGFDAAELVGKVHDKVEVVKP 69
XMreB2_zrk13           ---------------MAEKYGLGIDLGTANLLVFLEKKGIIFNEPSVIAFDRESGQIVAAGKDAHKMLGKVHDKISVIKP 65
XMreB3_zrk13           -------MIIVANKTNKKYIKIGVDLGTANLLVYVDGEGIIFNEPSVIAMEYASNDVIATGFNAARMIGRGHHGIKIVSP 73
IMreB1_HR1             -----------MGEKPKGTFNVGVDLGTSNLLIYVEGRGTIFDEPSYIAVDKASGKIVSVGLDAAELVGKVHDKVKVVKP 69
IMreB2_HR1             ---------------MAEKIGLGIDLGTANLLVYLEKKGIIFNEPSVIAFDRESGKIVAAGKDAHKMLGKVHDKISVIKP 65
IMreB3_HR1             ------MANDRKMKNNHKLIKIGIDLGTANLLVYVDGEGIIFNEPSVIAMEYETNDVIAVGFNAAKMIGRGHHGIKIVSP 74
IMreB1_HR2             ------MENNKLMKSKQKLIKIGIDLGTANLLVYVDGEGIIFNEPSVIAMDYDTSDVIAVGYNAAKMIGRGHHGIKIVSP 74
IMreB2_HR2             ---------------MADKIGLGIDLGTANLLVYLERKGIIFNEPSVIAFDRESGRIVAAGTDAHQMLGKVHSKISVVKP 65
IMreB3_HR2             -----------MSDKQKGKFNVGVDLGTSNLLIYVEGRGTVFNEPSIIAIDKATKKVVKVGKEAAELVGKVHDKIEVIKP 69
HMreB B1_SSD-17B       --------------MAGKGVKIGVDLGTVNTLVYINGQGVIYNEPSVIAFDRKTKKCIAVGKEAKDMVGKEHDHIDVIKP 66
HMreB A1_SSD-17B       --------------MASKKLGIGIDLGTTNLLVYVEGNGIIFNEPSVVAFDVETKDVIAGGTDAFSMIGKEHKKIKVSRP 66
HMreB B2_SSD-17B       --------------MAKKGYKIGVDLGTANTLVYINGHGLIYNEPSIVAFDKVTGKCIAAGNSASLMDGKQHERIRVVKP 66
HMreB A2_SSD-17B       --------------MAGKKLGIGIDLGTTNLLVYVQGHGIIFNEPSVVAFDVETGDVIAGGEDAFSMLGKTHKKIRVSRP 66
HMreB C2_SSD-17B       -------------MSNKEELKLGIDLGTVNTLVYANSRGIIFNEPTVIAFDKETDEVVAVGNEANDMVGKTHAKVRVVKP 67
HMreB A3_SSD-17B       -------------MANKKMLNIGIDLGTTNILVYVSGQGVIFNEPSVVAFDVDSGEVIAGGQAAYNMVGKTHGKIRVARP 67
HMreB B4_SSD-17B       ---------------MGQGYKIGVDLGTKNTIVYVGGKGIIYNEPTVVAFDQESKKPIAIGHSASEMLGKEHKKVKVVKP 65
HLPCO_02877_B5_bin.1   --------------MAKKSYRIGVDLGTANILVYITGSGVVYNEPSVVAFDNTTKECIAVGKRAFDMVGKEHKHIRIVKP 66
HLPCO_02873_B5_bin.1   --------------MAKKGYKIGVDLGTVNTLVYLNGQGLIFNEPSCVAFDRRTNDCIAVGQKAKDMLGREHSLIKVVKP 66
HLPCO_01615_B5_bin.1   --------------MKKSIYQIGIDLGTVNTLVYVSSRGIVYNEPTVVAFDSKTNKSFAIGHTAYDMLGKEHGKIKVVKP 66
HLPCO_02875_B5_bin.1   --------------MDKNNLKIGIDLGTANTLVYSSGRGVVFNEPTVVAFDKGTDKIIAIGHEAKEMLGKTHDKIRVVKP 66
HLPCO_02878_B5_bin.1   --------------MANKKLGIGIDLGTTNLLVYIEGHGIIFNEPSVVAFDIETGEVIAGGEDAFSMIGKTHKKIRVSRP 66
HLPCO_02874_B5_bin.1   --------------MPEKKIGIGIDLGTTNLLVYVTGHGIIFNEPSVVAFDIQTGEVIAGGEDAFSMVGKTHKKIRVSRP 66
B7766_RS04460_4B       --------------MAKKSHKIGVDLGTANILVYINGSGVVYNEPSVVAFDKGTNKCIAVGKRASEMLGKEHKHIRLAKP 66
B7766_RS04480_4B       --------------MVKKGYKIGVDLGTVNTLVYINGHGLIFNEPSCVAFDRRTNNCIAIGHNAYAMMGKEHHLVKISKP 66
B7766_RS04470_4B       --------------MTKNETKIGIDLGTANTLVYSSGRGVIFNEPSVVAFDKDTDQIIAIGSNAKSMLGKQHDKIRVVRP 66
B7766_RS04475_4B       ---------------MEKKLGIGIDLGTTNLLVYVQGNGIIFNEPSVVAFDIETGEVIAAGEDAFSMIGKTHKKIRVSRP 65
B7766_RS04455_4B       --------------MAIKKLGIGIDLGTTNLLVYYEGQGIIFNEPSVVAFDIETGEVIAGGNDAFSMIGKTHKKIQVSRP 66
BMreB_168              -------------MFGIGARDLGIDLGTANTLVFVKGKGIVVREPSVVALQTDTKSIVAVGNDAKNMIGRTPGNVVALRP 67

SMreB1_DSM21848        MVDGVITDIRATEAQLKYIFNR-LHVS--KQLKGSVMLLACPSVITELEKNALKKIATNLGADRVFVEEEVKMAALGGGV 147
SMreB2_DSM21848        LVDGVISDMDAAQDLLKHIFGR-LKMT--GIWKNSLVILACPSGVTELERSALKAIAKDMGASYVLVEEEVKLAALGAGI 148
SMreB3_DSM21848        LRNGAITDLRIAEEFIQHIGNR-AKVQ--DVWKGSIVLIACPKSVTELERRAMVEMCKHLGADLVQVEEDTLMAALGAGA 157
SMreB4_DSM21848        MVDGVITDIRATEAQLRYIFGK-LRIS--KQLKHSIMLLACPSVITELEKAALKKIAMNLGATKVFVEEEVKMAALGGGV 150
SMreB5_DSM21848        LVDGVIADMEAAQDLLKHVFSR-MKMM--NIWKNAVVLLACPSGVTELEREALKNVAQDMGADLVIIEEEAKMAAIGAGI 143
XMreB1_zrk13           LQGGVISDVSMIREILNFTFDQ-LFVN--SSQQINKLLICIPSEITETEKAAILQLGEELGIDDTKIDEEIKAAAIGSGV 146
XMreB2_zrk13           LKSGVISDMKAAKALLTYVLQE-VENLTEKDLTNTSCVMCCPSEVTKIERDIISELAMNMGISDVLIDEEIKAAALGANL 144
XMreB3_zrk13           LNQGVISDMDAAKKLIEIAVHK-AETI-DVNLQTSTLLICCPSEVTQIERDALIDLAHNLGVPDVFIEEEVKAGGIGAGL 151
IMreB1_HR1             LQGGVISDISMIRELLMFTFDK-LFVD--NTQQINKLLICIPSEITETEKAAILQLGAELGIEDTKIDEEIKAAAIGSGV 146
IMreB2_HR1             LRNGVISDMKAAKALLQYVLET-VENLTKKELTNTSCVMCCPSEVTTIERDIITELAVNMGISDVLIDEEIKAGALGANV 144
IMreB3_HR1             LNQGVISDMDAAKKLIEIAVRK-GEAI-DVNLRASTLLICCPSEVTQIERDAMIDLAHHLGVPDVFIEEEVKAGGIGAGL 152
IMreB1_HR2             LNQGVISDMEAAKKLIEIAIRK-AENS-DIKLKASTVLICCPSEVTQIERDAMIDLAHKLGVPDVFIEEEVKAGGIGAGL 152
IMreB2_HR2             LKNGVISDMKAAKALIAYVLNK-VENLTSKELSKTSCVICCPSEVTKIERDIMVDLALNTGISDVFIEEEIKSGALGANV 144
IMreB3_HR2             LNGGVIADMDMIRELLIFTFEK-LFVS--NLSKINRLLICIPSDITDTEKTAIMLLGQELGIEETYIDEEIKAAAIGGGI 146
HMreB B1_SSD-17B       LEGGVISDLEATKAYLQYVFEK-LEHI-NVKFKKSTLLICCPSEVTNIEKNAMGQLATQVGIRDVFIEEEIKAGAIGAGI 144
HMreB A1_SSD-17B       LRDGVISDMDAAKAMLKYVFER-VQNI--RDFHNATCLICCPSEVTQIEREAMRDLAIQMGIKDVFIEEEIKAGAIGAGI 143
HMreB B2_SSD-17B       LEGGVISDLDATKANLLYVFDK-LEHI-NVDFKKSTLLICCPSEVSTIERVAMKALASKMGINDVFIEQEVKAGAIGAGI 144
HMreB A2_SSD-17B       LRDGVISDMDAAKAMLRYVFKR-VQTI--RNFKDSKCLICCPSEVTQIEREAMRDLAIQMGIKDVFIEEEIKAGAIGAGI 143
HMreB C2_SSD-17B       LVSGVIADKNAAIRLIEYILNDFLEHYQSAKLKKSTVLICCHSDLSGVERNALKDIVQGFGINNVLVQEEVKAGAIGVGI 147
HMreB A3_SSD-17B       LREGVISDMEAAKALLKYIFLS-LKNR--KDLRNSYCVICCPSEVTHIERDAMRDLALQMGIGEVLIEEEIKAGAIGSGI 144
HMreB B4_SSD-17B       LDGGVISNLEATKIILSFIFNK-IQEDGNGDIKKSTLLICCPCDMTSTEREALEKLGTELNIKDVFVEQEIKAGAIGAGV 144
HLPCO_02877_B5_bin.1   LDGGVIADLDATKVYLQKVFEK-LENI-NVDLKNSTLLLCCPSQVTHIERVALIDLAKKIGVKDAFIEEELKAGAIGAGL 144
HLPCO_02873_B5_bin.1   LEGGVIADLDATKALLVYVFDK-LTSI-NIEFNRTTLLICCPSEVTQIERVALIALAKKLGVKDVFIEEEVKAGAIGAGI 144
HLPCO_01615_B5_bin.1   LEGGAIADLNATKVILKSILNK-IKIN-NIDFKKSTLLICCPSEMTTTERKALEKLGKELKINDVFVEQEIKAAAIGAGI 144
HLPCO_02875_B5_bin.1   IEQGVIADKDSAIQMLDHILKHHIMQFENFNPKKTTVLICCHSDLSMIERKALKDIIVNFGIQNVFVQEEVKAGAIGAGI 146
HLPCO_02878_B5_bin.1   LREGVISDMDAAKAMLKYVFQR-VQNV--RDFKHSKCLICCPSEVTQIEREAMRELAIQMGITDVFIEEEIKSGAIGSGI 143
HLPCO_02874_B5_bin.1   LREGVISDMDAAKAMLKYVFQR-VQNI--KNLKNAKCLICCPSEVTQIEREAMRELAVQMGINDVFIEEEIKSGAIGSGI 143
B7766_RS04460_4B       MEGGVIADLEATKVYLQYVFEK-LENI-SVDLKNSTILLCCPSEVSDIEKVALLELANKIGVGDAFVEQELKAGAIGAGI 144
B7766_RS04480_4B       LEGGVIADLDGAKELMKHVFTK-LVDI-NVDFKNSTLLICCPSEVSSIERAAIRDLSNNLGIKDVFVEEEVKAGAIGAGL 144
B7766_RS04470_4B       IEQGVVADKDATIKMLEYILNNHLINFENVKLKNALVLICCHSDLSGIERQALKDIMLNFGIQNVIVQEEVKAGAIGAGI 146
B7766_RS04475_4B       LREGVISDMDAAKALLRHVFER-VQNM--RDFKNSKCLICCPSEITQIEREAMKQLAIEMGINDVFIEEEIKAGAIGSGV 142
B7766_RS04455_4B       LREGVISDMDAAKALLKYVFQR-VQNM--RDFKNSKCLICCPSEVTQIEREAMRQLAIEMGINDVFIEEEIKSGAIGSGV 143
BMreB_168              MKDGVIADYETTATMMKYYINQAIKNK-GMFARKPYVMVCVPSGITAVEERAVIDATRQAGARDAYPIEEPFAAAIGANL 146
```

```
SMreB1_DSM21848          DIYKPNGNLVVDMGGGTTDVAVLSSGDIVLSKSVKVAGNYLNDEILKYVRSQYGLEIGIKTAEMIKIEIGSLAKYPDERK 227
SMreB2_DSM21848          NIGLAQGNLVIDIGGGTTDIAILSAGDIVKSKSVKVAGKHFDQEIQKYIRAEYNVLIGIRTAEQIKKDIGALVKIVNEKP 228
SMreB3_DSM21848          NIFAPKGTFILDIGGGKTSAGIISAGGIVVSKSIKIAGNYIDEEILKYIRAKHTISIGVVTAEQIKKQIGSLYKGKETKK 237
SMreB4_DSM21848          DIYKPTGNLVVDMGGGTTDIAVIASGDIVLSKSVKVAGNYLNDEMQKFIRSQYGLEVGSKTAEQIKIEIGSLAKYPDERK 230
SMreB5_DSM21848          NIDLPQGNLIIDIGGGTTDLAIISSGDIVVARSIKVAGNHFDDDIRKYIRSEYNIAIGQKTAEDVKKYIGSLVKYHNERA 223
XMreB1_zrk13             DIYTPSGHLVVDLGGGTTDFGVLSLGDVVLSKSIKVAGDYFDKQISDYVKEKHKLEIGPQTAEKAKIALASLTGDMPKDE 226
XMreB2_zrk13             DIFQSKGIMMVDIGGGTTDVGVLSFGDIVLSRTIRMAGNYIDTELAKLVKQNDKVEIGELTSENAKIELADLRKDAKLKV 224
XMreB3_zrk13             DIYDSVGSMVIDIGGGSTDIGVLSLGDIVVSESIRIAGNYLDQEIINYLQYEKGILIGKKTAQRIKEEIGTLRQEIVEEK 231
IMreB1_HR1               DIYTPSGHLVVDIGGGTTDFGVLSLGDVVLSKSIKVAGDYFDKQIMDYVKEEHKLEIGPQTAEKAKVALASLTGELPKDE 226
IMreB2_HR1               DIFKSKGVMIVDIGGGTTDVGVLSFGDIVLSRTIRMAGNFIDKELAKQVKQNEKVEIGELTSERCKMELADLRKDAKVKV 224
IMreB3_HR1               DIYDSKGSMVIDIGGGSTDIGVLSLGDIVVSESIRIAGNYLDQEIINYLQYQHGILIGKKTAQRIKEEIGTVRENLPGNV 232
IMreB1_HR2               DIYSSNGSMVIDIGGGSTDIGVLSLGDIVVSESIRVAGNHFDQEIINYLQYEHGLLIGKKTAQRIKEEIGTLREELTENK 232
IMreB2_HR2               DIFKSKGVMIVDIGGGTTDVGVLSFGDIVLSRTIRMAGNYVDTQITKYVKKDFKVEIGELTAERSKIELSDLRKDSSEII 224
IMreB3_HR2               DIYTPSGHLVVDIGGGTTDFGVLSLGDVVLSKSIKIAGDYFDKQIISYVKENHKLEIGLQTAERIKITLASLTGDLPINE 226
HMreB B1_SSD-17B         DIFAPQGAMIIDIGGGTSDIGVLALGDLVVSESTRIAGNYLDNQITKYVKMKYNMAIGNKTAETIKIKLGTLKETLEEEK 224
HMreB A1_SSD-17B         DIYSPSAAMVIDIGGGTTDVGVLSLGDVVLSHSIRVAGNYIDQEIIKYVHKHHNLIIGQKSAERAKIKVATLLEDEEEER 223
HMreB B2_SSD-17B         DIFAARGSMVIDIGGGSTDIGVLALGDLVVAESVRVAGNYIDRQIIKYVKMNHNMLIGFRTAEKIKVALGTVKEDLTDEK 224
HMreB A2_SSD-17B         DIYTPTAAMVIDIGGGTTDVGVLSLGDVVLSHSIRIAGNYIDQEIVKYVHKQHNLVIGHSSAEKAKKKIGTLLETDELQT 223
HMreB C2_SSD-17B         DIYSPTGSMIIDIGGGSTDVGVLALGDIVVSNSVKTAGNHLDAEIKKYIKMKRNFEIGMSTAERVKIALATVREDLVEEK 227
HMreB A3_SSD-17B         DIYSPEGSMVIDIGGGTTDVGVLSLGDVVLSQSIRVAGNYIDQIIVGYIHKSHNLVIGQKTAEKAKIETATLLDHVEFNS 224
HMreB B4_SSD-17B         DIFTAKGSLVIDIGGGSTDVGVLSLGDLVIAESIRKAGQYIDNEIIKYVKVHYGMIIGERTAEQIKINLGTLRKDIEEDK 224
HLPCO_02877_B5_bin.1     DIYESKGSMVVDIGGGTTDVGVLSLGDLVVSESCKVAGNYLDKEITKYVKFKYGIIIGASTAEMIKIKLGTLASELQEEK 224
HLPCO_02873_B5_bin.1     DIFAPQGSMVIDIGGGTTDIGVLSLGDLVVSESIRVAGNYIDNAIIKYVKMKYSMAIGSRTAENIKKSLGTVKQDLIEDK 224
HLPCO_01615_B5_bin.1     DIFTAKGSMIIDIGGGSTDIGVLSLGDLVVWDSIRIAGNSIDKEIMKYVKNKYGMLIGERTAEKIKMNIGTLRDDYNEDK 224
HLPCO_02875_B5_bin.1     DIYTPEGSMIIDIGGGSTDIGVLALGDIVVSNSIKVAGNHFDEEIRKYIKKKHNFEIGVLTAEEVKKALATLRDDLEEEK 226
HLPCO_02878_B5_bin.1     DIYHANAAMVIDIGGGTTDVGVLSLGDVVLSQSIRVAGNYIDQEIIKYVHKTHNLVIGQKTAERAKIEIATLLDE--EEK 221
HLPCO_02874_B5_bin.1     DIYTANAAMVIDIGGGTTDVGVLSLGDVVLSQSIRVAGNYIDQEIVKYVHKTHNLVIGQKSAEKAKIEIATLLDD--EER 221
B7766_RS04460_4B         DIFEASGSMVVDIGGGTTDIGVLSLGDLVISDSSKIAGNYLDKEISKYVKYKYGIIIGNNTSEQIKIQLGTLSKELSEDK 224
B7766_RS04480_4B         DIFAPIGSMIIDIGGGSTDIGVLALGDLVVSESIRVAGRYLDNQIIKYCKMRHGMAIGSRSAEKVKLEIGTLRRDLPNEK 224
B7766_RS04470_4B         DIYNPSGSMVIDIGGGSTDVGVLALGDIVVSKSIKTAGNAFDEEIRKFIKKKYNFEIGALSAEKVKISLATLRDDLPNEK 226
B7766_RS04475_4B         DIYTANATMVIDIGGGTTDVGVLSLGDVVLSQSIRVAGNYIDQEIVKYVHKTHNLVIGQRTAERVKMEISTLLDEEEEKF 222
B7766_RS04455_4B         DIYSANATMVVDIGGGTTDVGVLSLGDVVLSQSIRVAGNYIDQEIIKYIHKTHNLVIGQITAERVKIEVATLIDEEEEKF 223
BMreB_168                PVWEPTGSMVVDIGGGTTEVAIISLGGIVTSQSIRVAGDEMDDAIINYIRKTYNLMIGDRTAEAIKMEIGSAEAPEESDN 226

SMreB1_DSM21848          ------MKVYGRDVVSGLPREIELVPEEIREVLKVPISRIIDLTVQVLEETPPELAGDIFQNGITICGGGLIKGIATYF 301
SMreB2_DSM21848          ------IRAFGRDIITGLPREVMIKPEEIKNVLLAPFSRITDLLVEVLEETPPELAGDVIRNGITICGGGALIRGIVKYF 302
SMreB3_DSM21848          ------MVIFGRDVVTGMPKETEILDSEIRKLLISIFSSITQLVTDILESTPAELAGDAVMNGLLVSGGCAQISGLKEFL 311
SMreB4_DSM21848          ------MKVYGRDVVSGLPREIEVTPEEVREVLKVPVSRIIDLTVQVLEETPPELAGDIFKNGITIYGGGALIKGIDRYF 304
SMreB5_DSM21848          ------MQIYGRDIVSGLPKEAKISSDEIRNVLLNAFSKITDLVIELLENTPPELAGDIMRNGITICGGGALIRNIDKYF 297
XMreB1_zrk13             SGKPLTFDVMGRDLVSGLPKKVTLEASEIRTILLDAFEAVKATLIATLETTPPELAGDLVDNGIIVTGGGAKIKGIKDYI 306
XMreB2_zrk13             ------NRYAGRDIVRGIPKWVDISSTDVKQVLEPTYDEVVKLISAVLKDTPPELSADIFEHGIYLTGGGSLIKGVEEYI 298
XMreB3_zrk13             E-----TYANGRDIVTGLPRRITVTQTEVRDIFVEPFRAIANAVLKVLQNTPPELSADIIKSGMLVNGGCALIDGVDEFL 306
IMreB1_HR1               EGKAITFNAMGRDLVSGLPHMVVLKGKEIRKIMTDSFESIKATLIATLEATPPELAGDLVDNGIIVTGGGAQIKGIKEYI 306
IMreB2_HR1               ------NRYAGRDIVKGIPKWVDISSTDVNNVITPTYEEVVKLIAAVLKDTPPELSADIYQHGILLTGGGALIKGVEEFI 298
IMreB3_HR1               F-----TYANGRDIVSGLPRKIKVAQEDIREIFIEPFKSISNAILKVLQNTPPELSADIIQSGMLINGGCALVDGVDEFL 307
IMreB1_HR2               T-----TLAKGRDIVSGLPRKIEVSQEEVRAVLIEPFNNISTAILKVLQNTPAELSADIIVNGMLINGGCALIEGVDEFL 307
IMreB2_HR2               ------NRYAGRDIVKGIPKWVDISSKDVREVLVPVYEEVVKLISAVLKDTPPELSADIFQNGILLTGGGALIKGVEEFI 298
IMreB3_HR2               EGNPLTYSAMGRDVDSGLPQQVIIQAEEIREVLLSCFDTIRSTLISTLEATPPELAGDLVDNGILITGGGAQILGIRKFF 306
HMreB B1_SSD-17B         E-----YAFSGRNLKTGLPCKMTVKQSEIRDIFLRAFETITNTAKKVLQQSPPELAADIFNDGIIINGGGALIEGVQAYF 299
HMreB A1_SSD-17B         V-----ARIAGRDLVTGLPRTVNVTSQEIKTILQPIFQDITNVAYTVLEQTPPELAADIVDNGIIVDGGGALIPGVKEYF 298
HMreB B2_SSD-17B         E-----FTYAGRNLRTGIPCRHVIKESEVQRLTARAFESILNIAKKVLQQTPAELAADIFNDGVVINGGGALIGGVKEFF 299
HMreB A2_SSD-17B         ------YRVSGRDLVTGLPSSVELNSDEIRNILLPIFDEIRNVAYTVLEQTPPELAADIVDNGIVVDGGGALIPGIKEYF 297
HMreB C2_SSD-17B         E-----MVVSGRNLTNGLPSRITVKQSEIRDVLRKPFQTIINAILKVLEQTPPEIATDIIENGVILNGGGSQIDGLKEFI 302
HMreB A3_SSD-17B         E-----FRISGRDLVTGLPRAIVITSEEVRELLQPVFEEIGRLVMSVLEQTPPELAADIVDNGIVVNGGGSLIPGVKDYF 299
HMreB B4_SSD-17B         E-----YSFAGRSLENNLPRRKTIKQSEIRNILVRIFDSISTKVINVLQGTPPELAADILENGIVINGGGALIDGVEEYF 299
HLPCO_02877_B5_bin.1     E-----CVFAGRNLTNGLPCKMTIKQSEVREIFLKGFEVVVNTIKKVLQQTPPELASDIYQSEIMINGGGAMIDGVKEYL 299
HLPCO_02873_B5_bin.1     E-----FTYAGRNIVTGLPKRMVIKQSEVKALMIRALNSIANVARKVLEKTPPELSSDIFKDGIVLNGGGALIDGAKEYF 299
HLPCO_01615_B5_bin.1     A-----FKFAGRNIKTSLPDKKTIKQSEVRMVILKAIKAIASRALKILQDLEPELSSDIFERGIVINGGGALIDGIKEYF 299
HLPCO_02875_B5_bin.1     E-----ISVSGRDLKRGLPARITIKQSEVRNVLKRSFNAIASQILKVLEVTPPELAADIIETGVVLNGGGALIDGLKEYL 301
HLPCO_02878_B5_bin.1     F-----VKISGRDLVSGLPSATKLSSKEIRKLLLPIFDEIVNVVYSVLEQTPPELSADIVDNGIIVNGGGALIPGITEYI 296
HLPCO_02874_B5_bin.1     F-----VRVSGRDLVSGLPSATKISSKEIRKLLLPIFDEIVNVVYAVLERTPAELSADLVENGIVVNGGGALIPGVKEYI 296
B7766_RS04460_4B         E-----YTFAGRNLSNGLPCKVTMKQSEVKEIFVRAFESIVNTIRKVLQNTPPELSSDIFENGIVINGGGALIHGIKEFL 299
B7766_RS04480_4B         E-----YTVSGRNIKTGLPGRLVLKQSEIRDIFVKAFEGITITTFKVLQQTPPELAADIFENGIMINGGGALIDGIKEYF 299
B7766_RS04470_4B         E-----ITIAGRDLKSGLPSKITIRQTEIRDVLKNSFKTIANVILKVLEVTPPEIAADIIDNGVVLNGGGALIDGLDAYL 301
B7766_RS04475_4B         ------IRVSGRDLVSGLPSATKISSKEVRKILLPVFDEIVNVVYSVLEITPPELSADIVDNGIIVNGGGALIPGIKDYI 296
B7766_RS04455_4B         ------VKVSGRDLVSGLPSATKISSKEVRKIILPIFDEIINVVYSVLEVTPPELSADIVENGIIVNGGGANIPGVCEYI 297
BMreB_168                ------MEIRGRDLLTGLPKTIEITGKEISNALRDTVSTIVEAVKSTLEKTPPELAADIMDRGIVLTGGGALLRNLDKVI 300

SMreB1_DSM21848          EDTLQLPAKIGEQPLLAVINGTKKFES------DIYDIM-RQEHIKNKELDY------ 346
SMreB2_DSM21848          ESIFQLKVRAAQDPLMCVIDGAKTYEK------NLGAVIERIELLEAKEYKI------ 348
SMreB3_DSM21848          ESYFQIPVKIAKNPQTAVIDGCIAYEK------EIRDRL-----IEENKKNK------ 352
SMreB4_DSM21848          TDTLQLPSKVGEQPLLAVINGTKKFES------DIYDIL-RQEQMHTKELDY------ 349
SMreB5_DSM21848          FDIFQLPTRTASDPLMCVIEGTRAFEKVIRKRIENGYYN-----FNDKGLLAGIGKKK 350
XMreB1_zrk13             EQATNVEVHISPTPLTDVVDGTKKLLK-----INKNHYF--------GEF-------- 343
XMreB2_zrk13             SDRIKVPVKVVHNPLTCVAEGSKYLLK------NRGDYL-----VNPLKL-------- 337
XMreB3_zrk13             FNEIGLDVHIAKNPLTAIVEGTKVLLQ------NRGNYY-----VKPVD--------- 344
IMreB1_HR1               EEVTHVDVHVSHSPMTAVVEGTKRLLK-----MSKNHYF--------GEF-------- 343
```

2025/03/17 14:02 Sequence Manipulation Suite

```
IMreB2_HR1            SSRIKVPVKVVSNPLTCVAEGSKYLLK------NRGDYL-----VNPLKL-------- 337
IMreB3_HR1            FKEIGLDVHIAKNPLTAIVEGTKVLLQ------NRGNYY-----VKPVD--------- 345
IMreB1_HR2            EKKIGLDVHIAKNPLTAIVEGTKVLLQ------NRGNYY-----VKPVD--------- 345
IMreB2_HR2            SDRIKVPVKVVSNPLTCVAEGSKYLLK------NRGDYL-----VNPLQL-------- 337
IMreB3_HR2            EDITGTDVHISNGPLIAVVEGTKKLLK-----MDRKHFF--------GDY-------- 343
HMreB B1_SSD-17B      ENQLNLKIKIAENPLTAIVEGSKLLLK------NRGNYL-----VKPADV-------- 338
HMreB A1_SSD-17B      ESMLSLPVRITENPLTAVVQGTKVLLR------NKGSYI-----AQHKD--------- 336
HMreB B2_SSD-17B      ERELNLPVRVAENPLTSIVNGTKLLLS------NRGNYL-----IKPVE--------- 337
HMreB A2_SSD-17B      EEQINLPVRIAENSLTAVVEGTKVLLK------NRGSYL-----VNPTD--------- 335
HMreB C2_SSD-17B      ESIINIEVKLSPYPLTAIVKGTKELLK------NNGDYL-----VAPTD--------- 340
HMreB A3_SSD-17B      EELLNISVNIADKPLTAVVEGTKVLLQ------NKGSYL-----NNKHSRE------- 339
HMreB B4_SSD-17B      ETVTGITTYISKYPLTSIAEG-KTLLK------NRGNYL-----VKPID--------- 336
HLPCO_02877_B5_bin.1  EEALSLKVKIAENPLTSIVEGTRLLLN------NRGNYL-----VKPSDY-------- 338
HLPCO_02873_B5_bin.1  EDQLNLKVRVSENALTAIVDGSKYLLQ------NRGNYL-----IKPVD--------- 337
HLPCO_01615_B5_bin.1  ESITGLQVDIIKNPLTSIAEGTKYLLK------NRGNYL-----VKPLD--------- 337
HLPCO_02875_B5_bin.1  EELCQLNIRISPEPLTAIVKGTKDLLK------NKGDYL-----VAPSD--------- 339
HLPCO_02878_B5_bin.1  EEMVNLPVKVANNPLTAVVEGTKVLLK------NRGNYL-----INPAD--------- 334
HLPCO_02874_B5_bin.1  EEMVNLPVKISETPLTDVVEGTKILLK------NRGNYL-----INPQD--------- 334
B7766_RS04460_4B      EEELQLQVKISVNPLTSIVEGTKLLLN------NRGNYL-----VKPYDY-------- 338
B7766_RS04480_4B      EEAIGLKMKISLNPLTAIVDGTKYLLK------NRGNYL-----IKPLD--------- 337
B7766_RS04470_4B      EDLVQLKIRVSPDPLTSIIQGTKDLLK------NKGDYL-----VAPND--------- 339
B7766_RS04475_4B      EDMVHLPVKISESPLTDVVEGTKVLLK------NRGNYL-----INPTD--------- 334
B7766_RS04455_4B      EEMVNLPVRVVDSPLTAVVEGTKVLLK------NRGNYL-----INPTD--------- 335
BMreB_168             SEETKMPVLIAEDPLDCVAIGTGKALE------HI--HL-----FKGKTR-------- 337
```

**Fig. S1. Multiple alignment of amino acid sequences constructed by MUSCLE.**
ID is shown on the left of each sequence. See spread sheet in file “Data set S1” for the details. Multiple alignments are colored by Sequence Manipulation Suite. The area marked by a purple box is the membrane binding site previously reported (*24*). However, this feature was not found in *Haloplasma contractile* SSD-17B sequences. These sequences were trimmed with the automatic1 option of trimAl and a phylogenetic tree was constructed in Fig. 1A.

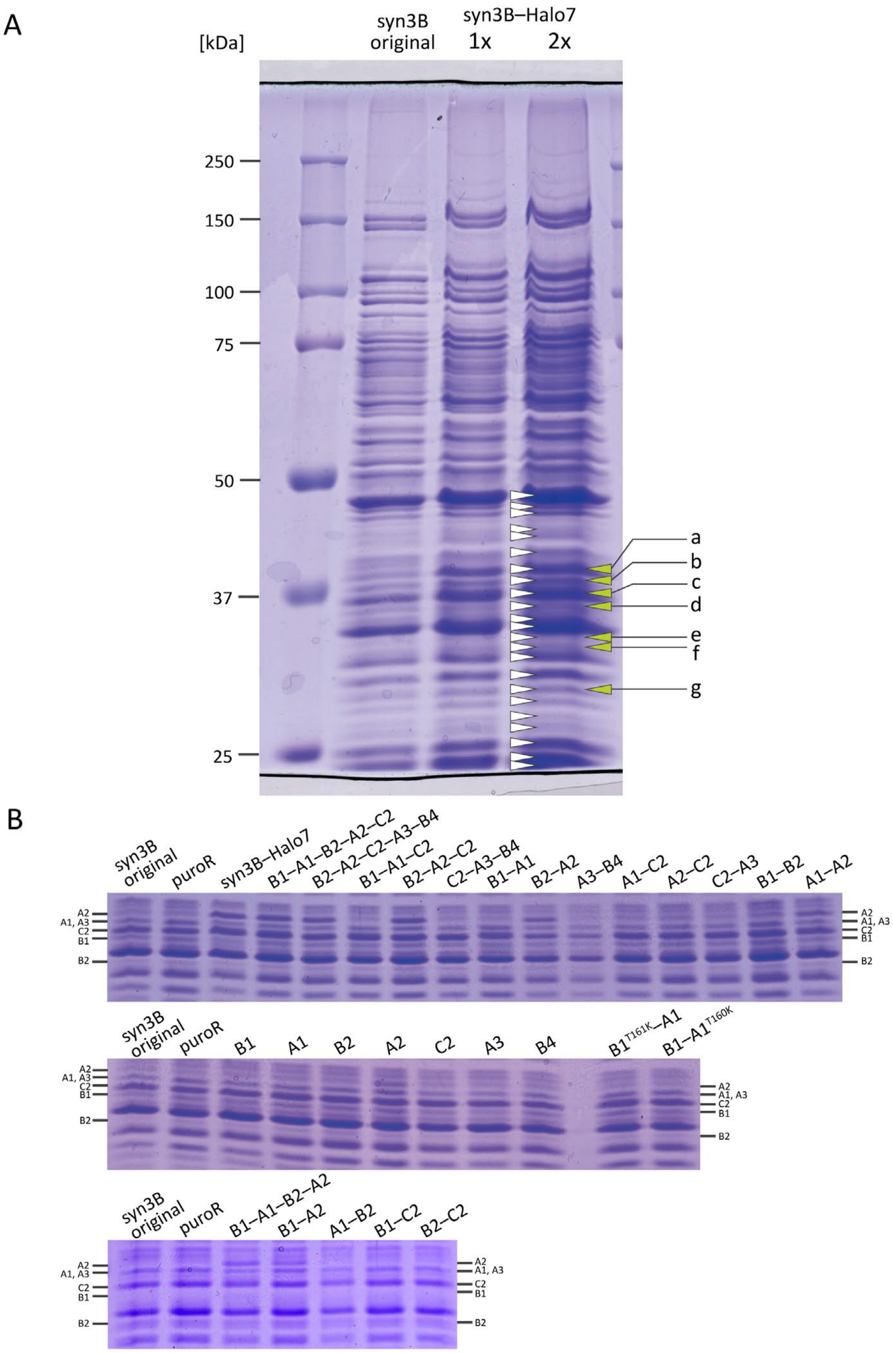


**Fig. S2. Protein profile of syn3B cell expressing *H*MreBs.** The cell lysates were analyzed by SDS-10% PAGE stained by Coomassie Brilliant Blue. (A) syn3B original and syn3B–Halo7. syn3B–Halo7 was analyzed with two different subjection amounts. Twenty-three bands marked by white triangles were analyzed by PMF based on MALDI-TOF mass spectrometry. The bands marked by green triangles were shown to include *H*MreBs. (B) syn3B cell expressing various combinations of *H*MreBs.

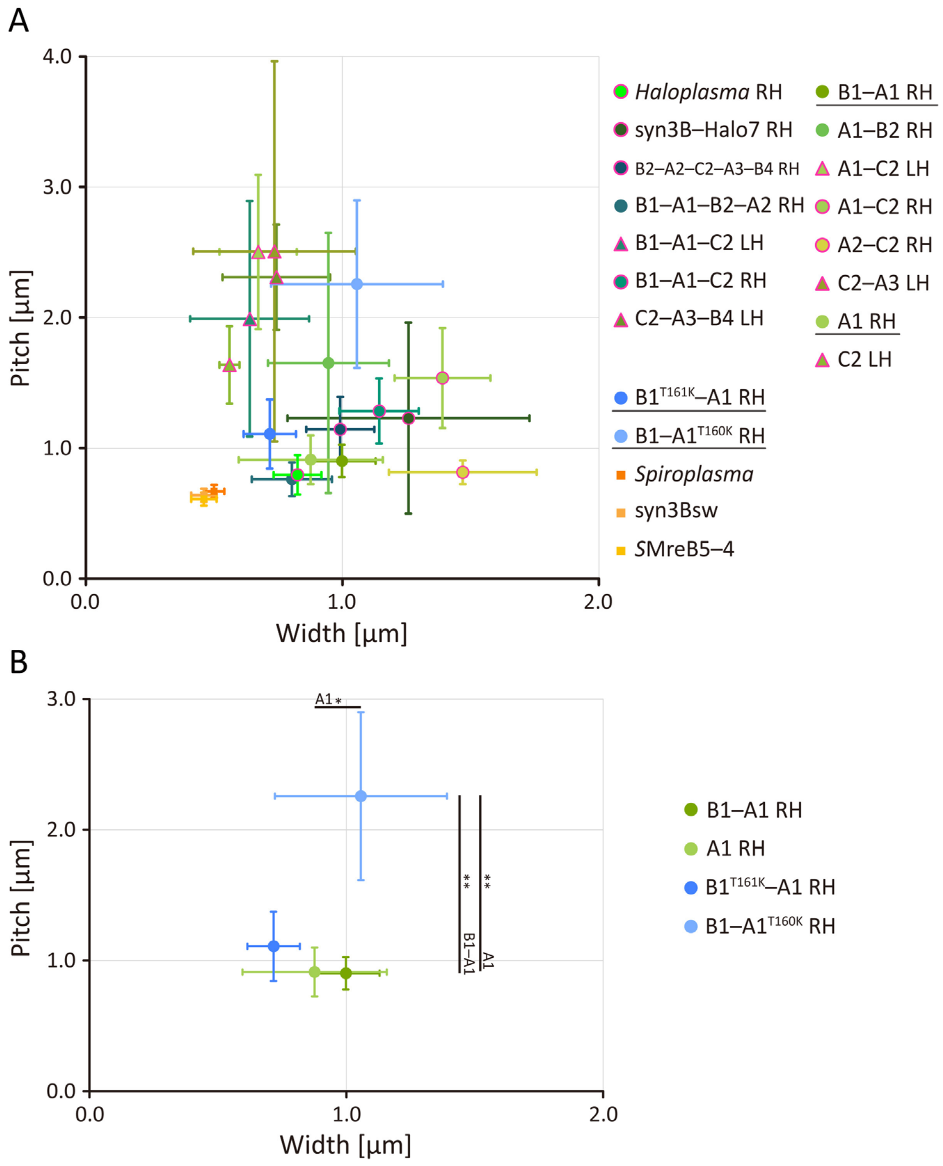


**Fig. S3. Dimensions of helical cells.**
(A) Summary of cell dimensions and handedness. Cell handedness is presented by circle, triangle, and square symbols for right, left, and mixed handedness, respectively. Involvement of *H*MreB C2 in protein expression is shown by magenta outline. At least three cells were analyzed for each strain. Data from *Spiroplasma eriocheiris*, syn3Bsw, and syn3B expressing *S*MreB5–4 (*S*MreB5–4) were copied from a previous paper (*11*), where helical handedness was not analyzed. (B) Influence of ATPase reduction of *H*MreB B1 and A1 on cell dimensions. Data from strains underlined in (A) are plotted here. The black bars show differences of B1–A1$^{T160K}$ strain from related strains with evaluation by Student's *t*-test (** $p < 0.01$, * $p < 0.05$).

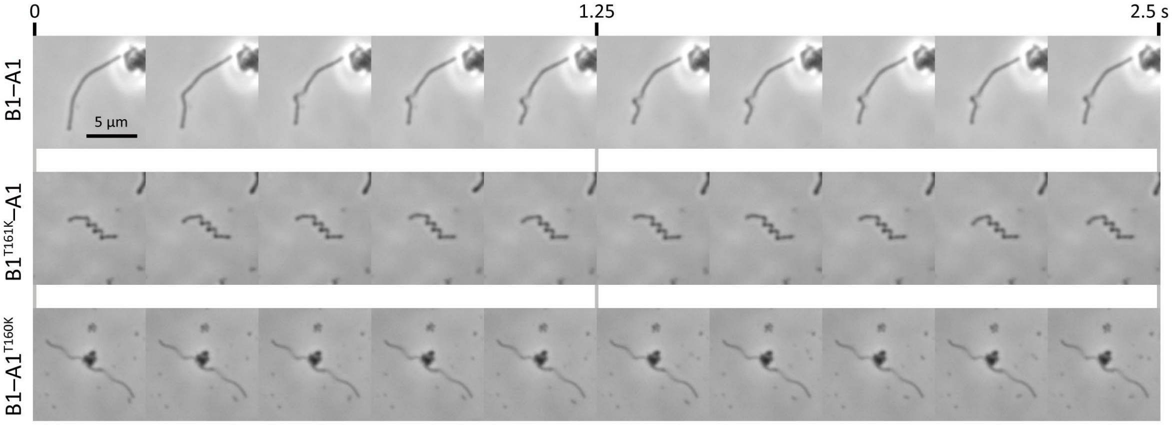


**Fig. S4. Cells expressing *H*MreB B1 and A1 with a mutation reducing ATPase activity.** Consecutive video images for 2.5 s. Strains are shown in the left.

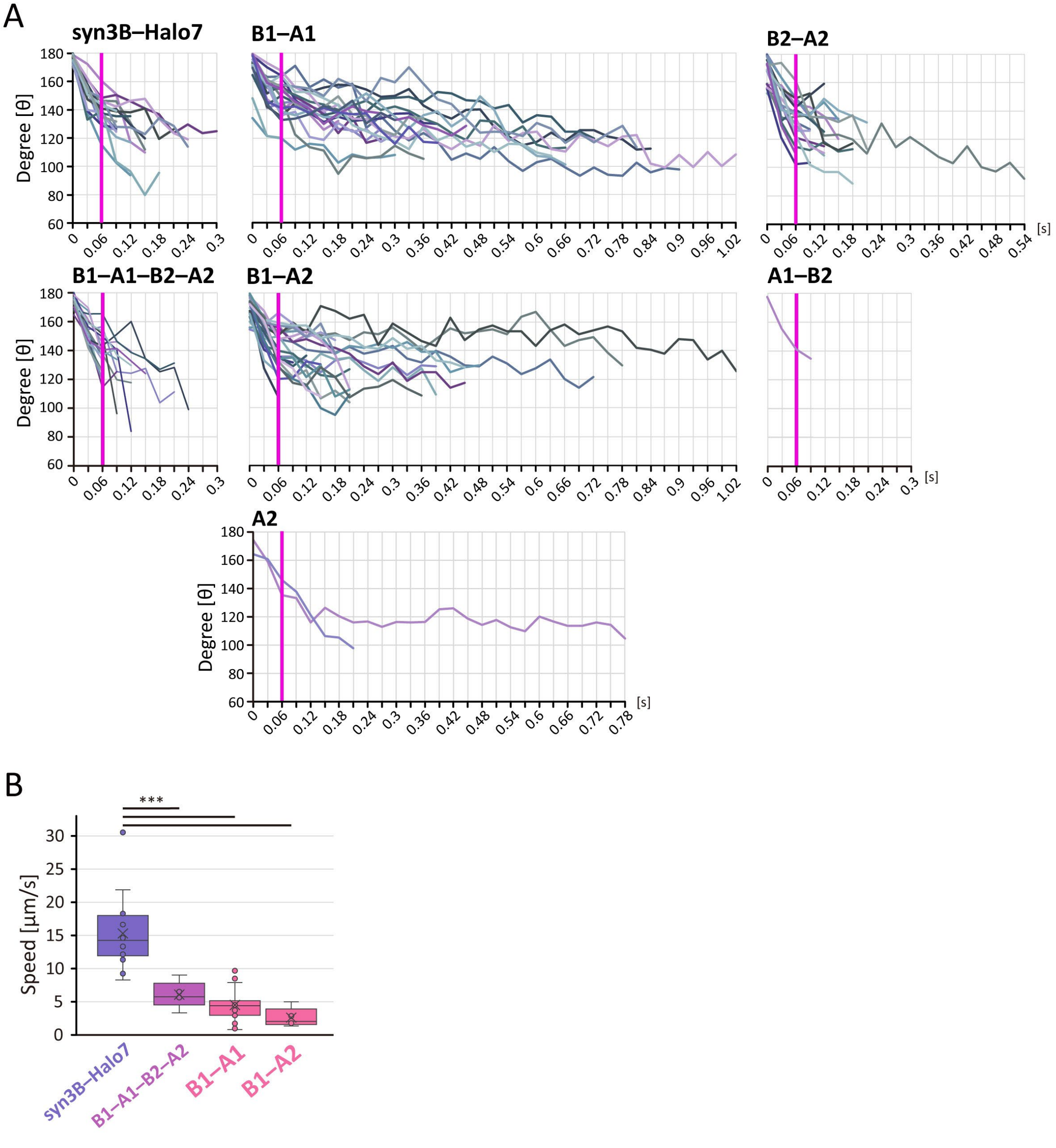


**Fig. S5. Comparison of traveling speeds.**
(A) Change in curving angle in type I movement, curve formation. The speed was measured for the initial 0.06 s. (B) Loop traveling speeds. Cells from syn3B–Halo7 (n = 20), *H*MreBs B1–A1–B2–A2 (n = 20), B1–A1 (n = 6), and B1–A2 (n = 6) strains were analyzed. Bars present *p*-value supported by Tukey-Kramer HSD test (*** $p < 0.002$).

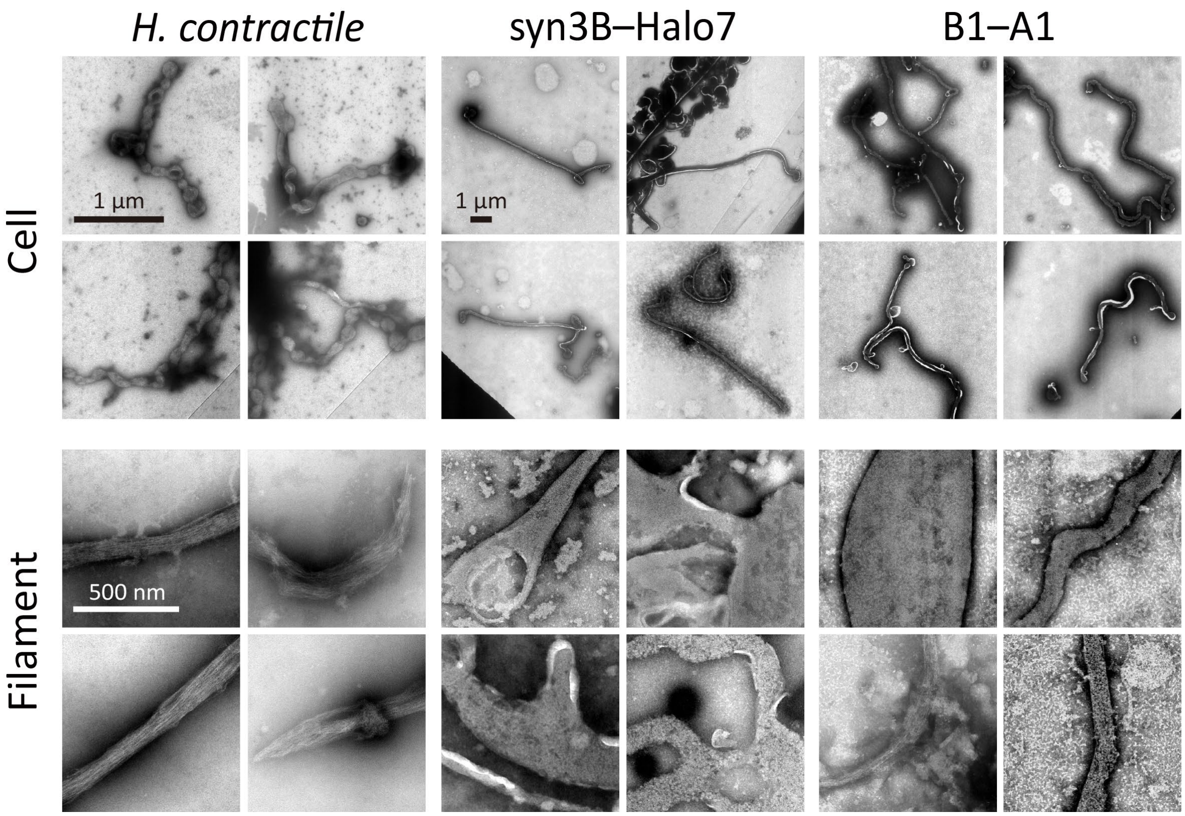


**Fig. S6. Negative-staining EM images.**
Cell (upper) and filament (lower) images obtained by negative-staining electron microscopy. The species or strain names are shown on the top.

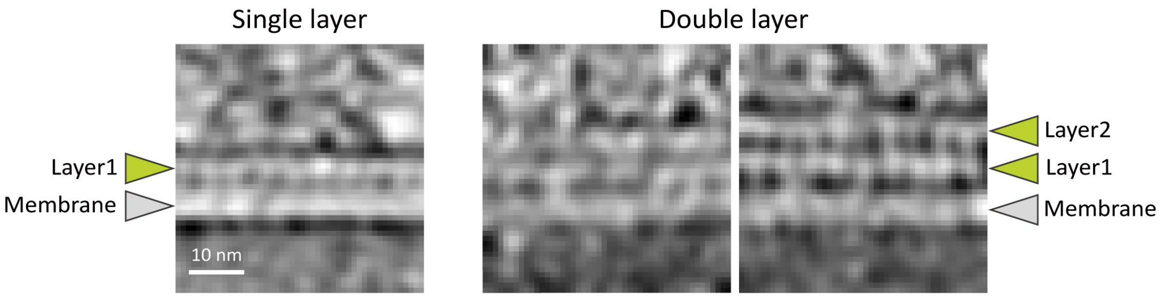


**Fig. S7. Double layer of filaments.**
Double layered filaments were observed in some positions as shown by smooth tomographic slices (right panel). A single layered filament image identical with that of Fig. 5D (left panel) is shown for comparison.

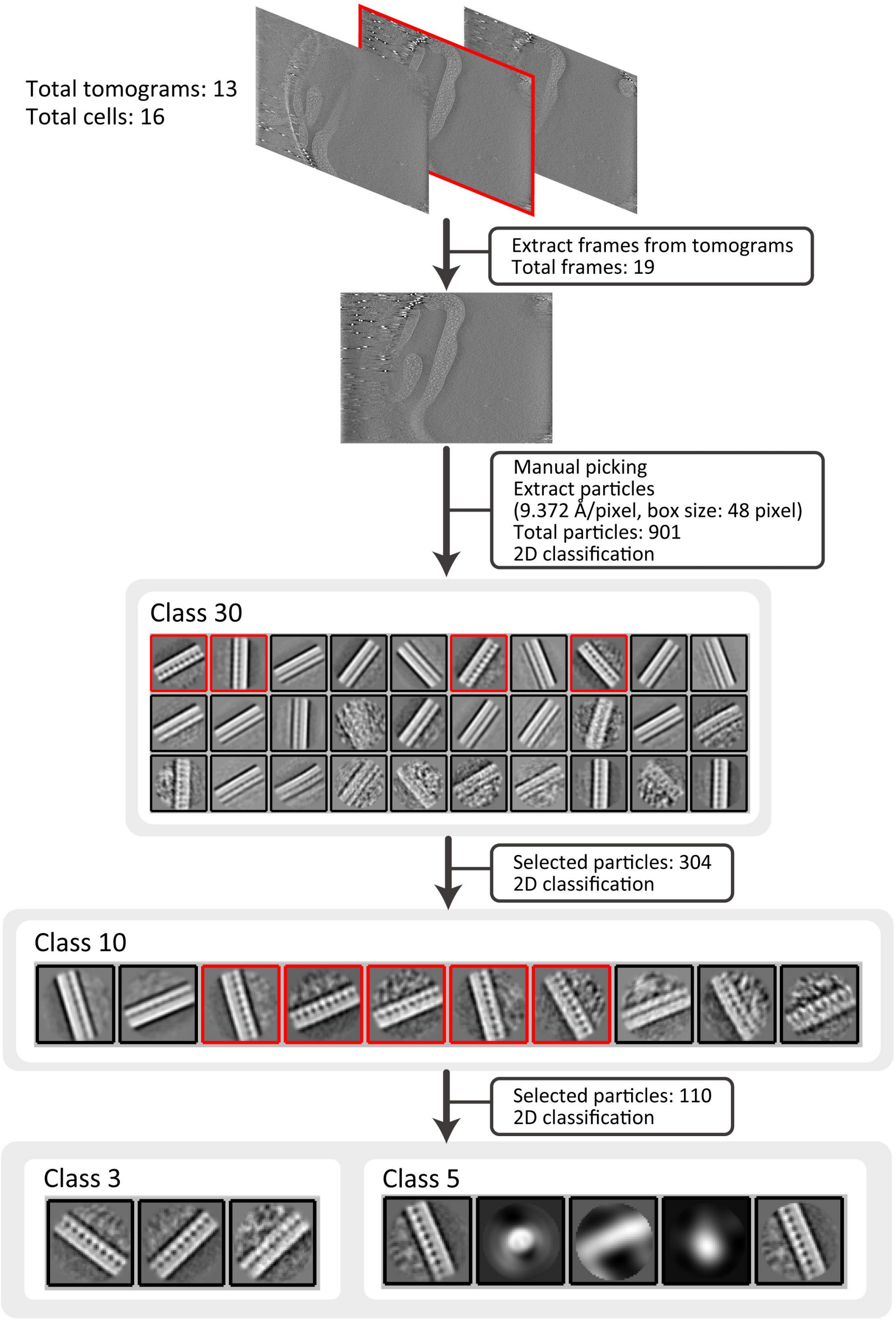


**Fig. S8. Analysis by cryo-electron tomography including two-dimensional averaging.**

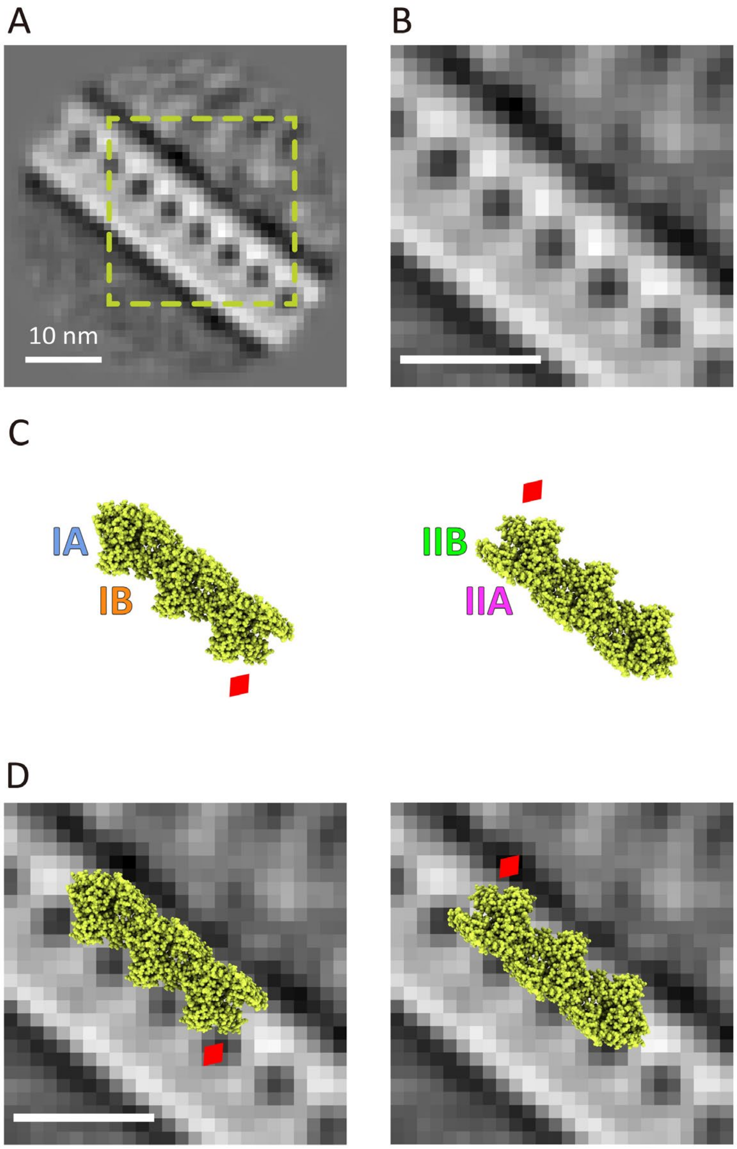


**Fig. S9. Overlay of the averaged image and *H*MreB A1.**
(A) Averaged image of *H*MreBs B1–A1 in syn3B obtained by cryo-ET. (B) Magnified image of boxed area in panel (A). (C) Filament structure of *H*MreB A1 predicted by AlphaFold3. The atomic diameter is modified from default to −0.3. The left image is rotated 180 degrees on a plane to be the right one. Red diamond marks a position of the filament. (D) The predicted structure shown in (C) was overlaid onto the image of (B) with different alignments.

### Fig. S10. DNA constructs used in this study.

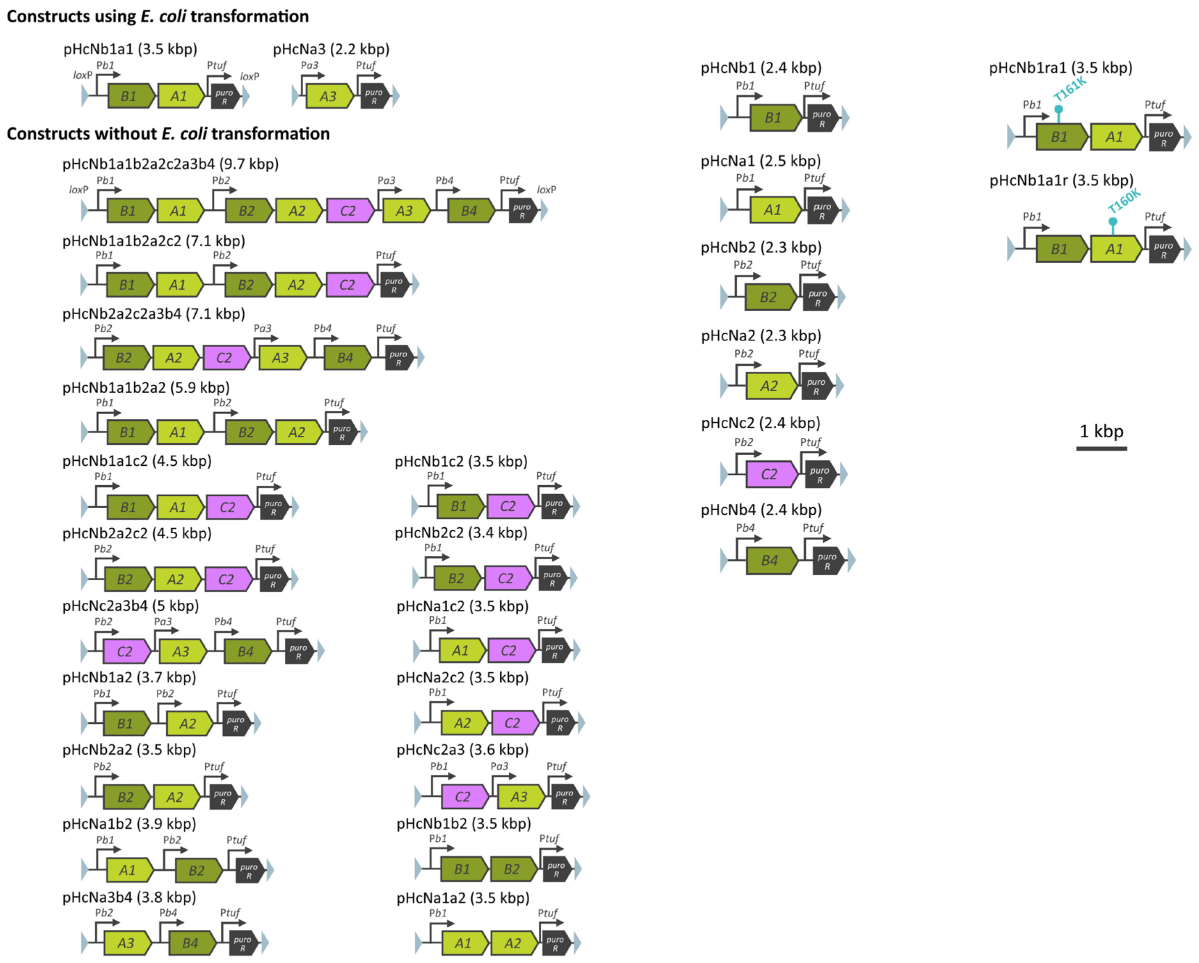


DNA constructs containing *hmreB* genes from *H. contractile* and *puroR* (a puromycin resistance gene). The promoter and *lox*P sites are shown by black arrow and light blue triangles, respectively. P** is the promoter predicted upstream of *hmreB* ** by the SAPPHIRE program.

**Table S1. Proteins identified by PMF for bands shown in Fig. S2.**

| Protein band | Database | Gene ID | Annotation | Mass (kDa) | Score | Coverage (%) |
|---|---|---|---|---|---|---|
| a | JCVI-syn3B | JOY36_00575 | acetate kinase protein | 44.4 | 88 | 34 |
| a | *H. contractile* | HLPCO_003151 | *H*MreB A2 | 36.5 | 93 | 44 |
| a | *H. contractile* | HLPCO_002796 | *H*MreB A1 | 36.6 | 28 | 15 |
| b | JCVI-syn3B | JOY36_01535 | 30S ribosomal protein S2 | 32.7 | 62 | 42 |
| b | *H. contractile* | HLPCO_002796 | *H*MreB A1 | 36.6 | 56 | 36 |
| b | *H. contractile* | HLPCO_003151 | *H*MreB A2 | 36.5 | 36 | 22 |
| b | *H. contractile* | HLPCO_003147 | *H*MreB A3 | 36.7 | 28 | 26 |
| c | JCVI-syn3B | JOY36_01630 | type I glyceraldehyde-3-phosphate dehydrogenase protein | 37.2 | 137 | 63 |
| c | JCVI-syn3B | JOY36_02335 | ribose-phosphate diphosphokinase protein | 38.6 | 69 | 40 |
| c | *H. contractile* | HLPCO_003150 | *H*MreB C2 | 36.9 | 96 | 60 |
| c | *H. contractile* | HLPCO_002796 | *H*MreB A1 | 36.6 | 30 | 23 |
| d | JCVI-syn3B | JOY36_01630 | type I glyceraldehyde-3-phosphate dehydrogenase protein | 37.2 | 99 | 46 |
| d | JCVI-syn3B | JOY36_01775 | DNA-directed RNA polymerase subunit alpha protein | 35.0 | 60 | 32 |
| d | JCVI-syn3B | JOY36_01225 | HD domain-containing protein | 37.5 | 56 | 39 |
| d | *H. contractile* | HLPCO_002797 | *H*MreB B1 | 36.9 | 33 | 32 |
| e | JCVI-syn3B | JOY36_01295 | L-lactate dehydrogenase protein | 34.8 | 72 | 46 |
| e | JCVI-syn3B | JOY36_00545 | 6-phosphofructokinase protein | 35.3 | 46 | 24 |
| e | *H. contractile* | HLPCO_003152 | *H*MreB B2 | 36.6 | 47 | 35 |
| f | JCVI-syn3B | JOY36_01820 | 30S ribosomal protein S5 protein | 27.8 | 57 | 49 |
| f | *H. contractile* | HLPCO_003152 | *H*MreB B2 | 36.6 | 39 | 32 |
| g | JCVI-syn3B | JOY36_00195 | Cof-type HAD-IIB family hydrolase protein | 32.4 | 78 | 50 |
| g | JCVI-syn3B | JOY36_01680 | DegV family protein | 31.8 | 50 | 42 |
| g | JCVI-syn3B | JOY36_02385 | transcription termination/antitermination protein NusG protein | 24.1 | 49 | 38 |
| g | *H. contractile* | HLPCO_003150 | *H*MreB C2 | 36.9 | 16 | 10 |

Score was provided by Mascot search version 2.5.1. Protein scores of Mascot greater than 39 or 47 are significant for JCVI-syn3B (taxonomy ID: 2806337) or *H. contractile* (taxonomy ID: 1033810), respectively.

### Table S2. DNA primers used in this study.

| Name | Sequence | Purpose |
|---|---|---|
| LP_inverse_F | AGGACTGAGCTAGCTGTCAAAGATC | pHcNb1a1b2a2c2a3b4, pHcNb1a1b2a2c2, pHcNb2a2c2a3b4, pHcNb1a1b2a2, pHcNb2a2c2, pHcNb1a1, pHcNb1a2, pHcNb2a2, pHcNb1c2, pHcNb2c2, pHcNb1b2, pHcNb2, pHcNa3, pHcNb4, pHcNb1ra1, pHcNb1a1r |
| LP_inverse_R | GAACTATATAAATAACTCGCATATTG | pHcNb1a1b2a2c2a3b4, pHcNb1a1b2a2c2, pHcNb2a2c2a3b4, pHcNb1a1b2a2, pHcNb1a1c2, pHcNb2a2c2, pHcNc2a3b4, pHcNb1a1, pHcNb1a2, pHcNa1b2, pHcNb2a2, pHcNa3b4, pHcNa1c2, pHcNa2c2, pHcNc2a3, pHcNb1c2, pHcNb2c2, pHcNb1b2, pHcNa1a2, pHcNb1, pHcNa1, pHcNb2, pHcNa2 pHcNc2, pHcNa3, pHcNb4, pHcNb1ra1, pHcNb1a1r |
| LP-H1-F | GCGAGTTATTTATATAGTTCTAGGGCGGTACCaaaattaaaaaagc | pHcNb1a1b2a2c2a3b4, pHcNb2a2c2a3b4, pHcNc2a3b4, pHcNa3b4, pHcNb4 |
| LP-H1-R | TTGACAGCTAGCTCAGTCCTGAATTCaaactcatctccttag | pHcNb4 |
| LP-H2-F | GCGAGTTATTTATATAGTTCttaattaaccgacagcccctt | pHcNb1a1, pHcNb1ra1, pHcNb1a1r |
| LP-H2-R | TTGACAGCTAGCTCAGTCCTtgctgggtagttttttatg | pHcNb1a1b2a2c2a3b4, pHcNb1a1b2a2c2, pHcNb1a1b2a2, pHcNb1a1c2, pHcNb1a1, pHcNb1a2, pHcNb1c2, pHcNb1b2, pHcNb1, pHcNa1, pHcNb1ra1, pHcNb1a1r |
| LP-H2-short-R | TAGCTCAGTCCTtgctgggtagttttttatg | pHcNb1a1b2a2c2a3b4, pHcNb1ra1, pHcNb1a1r |
| LP-H3-F | GCGAGTTATTTATATAGTTCGGTACCAGCACACATTAAAAAGG | pHcNa3 |
| LP-H3-2209-R | TTGACAGCTAGCTCAGTCCTGTAAGACAAGACACACGAAGACA | pHcNa3 |
| LP-H4-F | GCGAGTTATTTATATAGTTCTGTAATGAAAAAGTCGTCATAAAG | pHcNb1a1c2, pHcNb2a2c2, pHcNa1c2, pHcNb1c2, pHcNb2c2, pHcNa2c2, pHcNc2 |
| LP-H4-short-F | GCGAGTTATTTATATAGTTCTGTAATGAAAAAGTCGTC | pHcNb1a1b2a2c2 |
| LP-H4-R | TTGACAGCTAGCTCAGTCCTCTAAAAATGCCAATTCCCGACAG | pHcNb2a2c2a3b4, pHcNb2a2c2, pHcNb2a2, pHcNb2c2, pHcNb2 |
| LP-hm1-F | GCGAGTTATTTATATAGTTCattcaaatcctcctcagattta | pHcNb1 |
| LP-hm2-F | GCGAGTTATTTATATAGTTCttatcaaatatagcgca | pHcNa1 |
| LP-hm3-F | GCGAGTTATTTATATAGTTCttgaatttcctccctttaga | pHcNa1b2, pHcNb1b2, pHcNb2 |
| LP-hm4-F | GCGAGTTATTTATATAGTTCATTAGTCAGTAGGATTTAC | pHcNb1a1b2a2, pHcNb1a2, pHcNa1a2, pHcNa2 |
| LP-hm6-F | GCGAGTTATTTATATAGTTCAGCACACATTAAAAAGG | pHcNc2a3 |
| LP-hm6-2308-R | GCTCAGTCCTGTAAGACACGAAGACAGAGTAACAA | pHcNb1a1b2a2c2a3b4, pHcNb2a2c2a3b4 |
| LP-hmreB3-4-F | GCGAGTTATTTATATAGTTCGTTTACTGTACCTAGGTCTATCCC | pHcNb2a2 |
| LP_inverse_Phm1_F | tttttcaccttctctaagtata | pHcNa1b2, pHcNa1c2, pHcNa2c2, pHcNc2a3, pHcNa1a2, pHcNa1 |
| LP_inverse_Phm3_F | CCTATCACCTCTATAATTT | pHcNc2a3b4, pHcNa3b4, pHcNa2, pHcNc2 |
| cn-hm23-F | CGGGAATTGGCATTTTTAGagtagttatattttatcaa | pHcNb1a1b2a2c2a3b4 |
| cn-hm23-R | tataactactCTAAAAATGCCAATTCCCGACAGCC | pHcNb1a1b2a2c2a3b4 |
| cn-hm56-F | CGTGTCTTACTGTAATGAAAAAGTCGTCATAAAGA | pHcNb1a1b2a2c2a3b4, pHcNb2a2c2a3b4 |
| cn-hm56-R | TTTCATTACAGTAAGACACGAAGACAGAGTAACAA | pHcNb1a1b2a2c2a3b4, pHcNb2a2c2a3b4 |
| cn-hm67-F | actaaggagatgagtttAGCACACATTAAAAAGGCCCCTT | pHcNb1a1b2a2c2a3b4, pHcNb2a2c2a3b4 |
| cn-hm67-R | TAATGTGTGCTaaactcatctccttagtaatgatt | pHcNb1a1b2a2c2a3b4, pHcNb2a2c2a3b4 |
| cn-hm25-F | CCTCATAAGTATCttagtctttatgttgagctat | pHcNb1a1c2, pHcNa1c2 |
| cn-hm25-R | ctcaacataaagactaaGATACTTATGAGGTGAGAAA | pHcNb1a1c2, pHcNa1c2 |
| cn-hm13-F | GCCATattcaaatcctcctcagatttattttatacatcag | pHcNb1b2 |
| cn-hm13-R | ctgaggaggatttgaatATGGCAAAGAAAGGATATAA | pHcNb1b2 |
| cn-hm24-F | TCCCTTTAGAATATttagtctttatgttgagctat | pHcNa1a2 |
| cn-hm24-R | tcaacataaagactaaATATTCTAAAGGGAGGA | pHcNa1a2 |
| cn-hm2-Phm1-R | tacttagagaaggtgaaaaaatggcaagcaagaaattagg | pHcNa1c2, pHcNa1a2, pHcNa1 |
| cn-hm4-Phm1-R | tacttagagaaggtgaaaaaATGGCCGGTAAAAAATTAGG | pHcNa2c2 |
| cn-hm5-Phm1-R | tatacttagagaaggtgaaaaaATGTCAAATAAAGAAGAATT | pHcNc2a3 |
| cn-hm4-Phm3-R | AAAATTATAGAGGTGATAGGATGGCCGGTAAAAAATTAGG | pHcNa2 |
| cn-hm5-Phm3-R | AAAATTATAGAGGTGATAGGATGTCAAATAAAGAAGAATT | pHcNc2a3b4, pHcNc2 |
| cn-hm6-Phm3-R | AAAATTATAGAGGTGATAGGGAATGATAAAGTGGCGAACA | pHcNa3b4 |
| cn-hm14-F | CGGGAATTGGCATTTTTAGattcaaatcctcctcag | pHcNb1a2 |
| cn-hm14-R | aggatttgaatCTAAAAATGCCAATTCCCGAC | pHcN1b1a2 |
| cn-hm15-F | GTTGCATCATATGTattcaaatcctcctcagatt | pHcNb1c2 |
| cn-hm15-R | aggatttgaatACATATGATGCAACTATTTCTAG | pHcNb1c2 |
| cn-hm14-R | aggatttgaatCTAAAAATGCCAATTCCCGAC | pHcNb1a2 |
| cn-hm15-F | GTTGCATCATATGTattcaaatcctcctcagatt | pHcNb1c2 |
| cn-hm15-R | aggatttgaatACATATGATGCAACTATTTCTAG | pHcNb1c2 |
| cn-hm35-F | GTTGCATCATATGTCCTCCCTTTAGAATATA | pHcNb2c2 |
| cn-hm35-R | ATATTCTAAAGGGAGGACATATGATGCAACTATTTCTAG | pHcNb2c2 |

| Name | Sequence | Purpose |
|---|---|---|
| 1r2_T161K_CtoA_F | tatatcagattttccaccaccgatat | pHcNb1ra1 |
| 1r2_T161K_CtoA_R | tggtggaaaatctgatataggtgttttag | pHcNb1ra1 |
| 12r_T160K_TCtoAA_F | aacatcagtttttcctccaccaatatc | pHcNb1a1r |
| 12r_T160K_TCtoAA_R | gtggaggaaaaactgatgttggagttttatc | pHcNb1a1r |
| coloP-puro-F | ggagtagtccaaacagcaacagca | PCR after recombination by NEBuilder HiFi DNA assembly® |
| seq_term_R | GGTGAAAACCTCTGACACATG | PCR after recombination by NEBuilder HiFi DNA assembly® |
| syn3B_junc_F | TATGTGATAATGCCAATCGCTAAG | colony PCR |
| syn3B_junc_R | GTAAATTCCCAAATTATTCCATCA | colony PCR |

**Supplementary text (Details of biophysical modeling)**
In this section, we briefly describe the details of the biomechanical modeling and the numerical method of our mathematical model.

**Elastic energy and the modeling of active shape changes**
Our bacterial cell body is sufficiently thin because the diameter of the cell, *a*, is typically much smaller than its contour length *L*. In our biophysical model, the bacterial cell body is thus regarded as a uniform elastic rod of length *L* and isotropic cross-section of diameter *a* (Fig. S11).

To describe the centerline configuration of the rod, we assign an orthogonal director frame $(\hat{\mathbf{d}}_1, \hat{\mathbf{d}}_2, \hat{\mathbf{d}}_3)$ at each point of the centerline *s*, where *s* is the arclength along the rod centerline measured at one end of the rod (at which $s = 0$). See Fig. S11. Here, $\hat{\mathbf{d}}_3 = \mathbf{r}'$ is, by definition, the unit tangent vector, where $()'$ denotes the derivative with respect to *s*, while $\hat{\mathbf{d}}_1$ and $\hat{\mathbf{d}}_2$ lie on the rod's cross-sectional plane. The rod configuration is determined by specifying how the director frame rotates as it moves along the centerline, $\hat{\mathbf{d}}_a' = \mathbf{\Omega} \times \hat{\mathbf{d}}_a$, where $a = 1,2,3$. The orientation of the vector $\mathbf{\Omega} = \Omega_a \hat{\mathbf{d}}_a$ sets the rotational axis at *s*, and $\Omega_a$ gives the rate of rotation, i.e., curvature, about $\hat{\mathbf{d}}_a$.

We assume that an equilibrium configuration of a cell can be fully determined as the energy minimizing state of an elastic rod with an intrinsic curvature $\kappa$ and/or an intrinsic torsion $\tau$. Within the framework of the Hookean elasticity, the bending and twisting elastic energy of an inextensible rod can be given by

$$E_{\mathrm{rod}} = \int_0^L \left[\frac{A}{2}\Omega_1^2 + \frac{A}{2}(\Omega_2 - \kappa)^2 + \frac{C}{2}(\Omega_3 - \tau)^2\right] ds, \tag{1}$$

where *A* and *C* are, respectively, the bending and twisting rigidities.

In our coarse-grained mechanical model, we assume that the cell's active shape dynamics is generated by the temporal and spatial changes of the preferred curvature $\kappa(s,t)$ or twist $\tau(s,t)$ in Eq. (1), as done in the previous studies [18, 65-68].

For a coiling *Haloplasma*, we assume that the preferred twist is uniform, i.e., $\tau$ is constant along the cell, while the preferred curvature changes according to the one-dimensional propagating wave with the constant phase speed $v_b (> 0)$, i.e., $\kappa(s,t) = \kappa_0 \psi(s - v_b t)$, where

$$\psi(s) = \frac{1}{2}\left[\tanh\left(\frac{s}{\xi}\right) - \tanh\left(\frac{s - D}{\xi}\right)\right]. \tag{2}$$

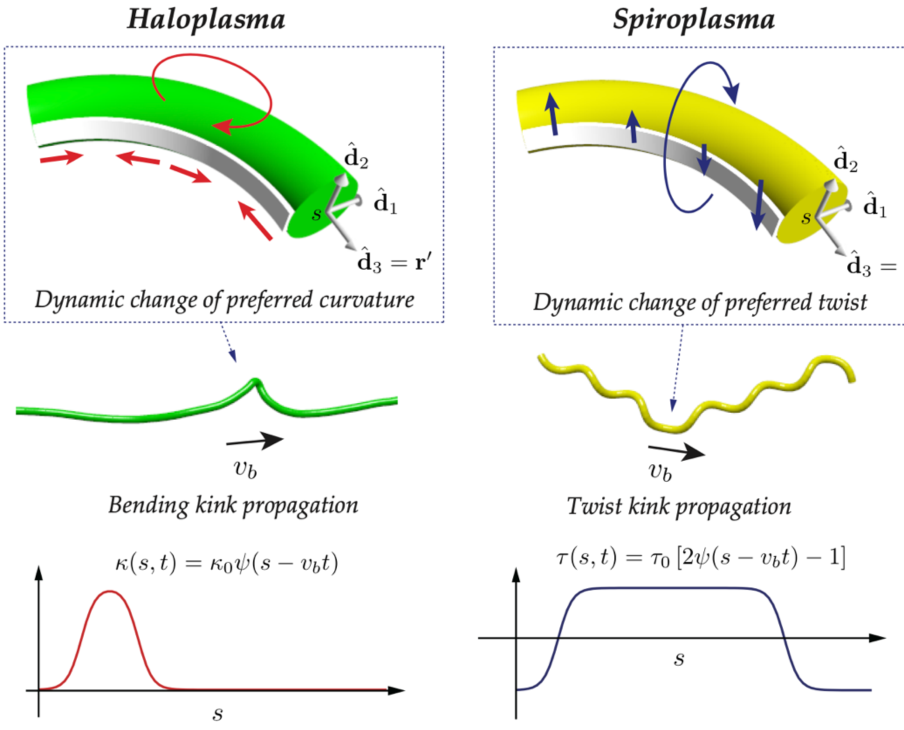


Fig. S11: Schematic picture of an elastic rod model of a filamentous *Haloplasma* (left) or *Spiroplasma* (right) cell. (Upper) Geometry of our rod model and the definition of the director frame at each cross section of the rod. A contractile MreB ribbon (white) is underlying a dynamic preferred curvature $\kappa(s,t)$ for *Haloplasma* (left) or a dynamic preferred twist $\tau(s,t)$ for *Spiroplasma* (right) postulated in our elastic model. (Middle) Overview of a filamentous cell with a localized kink region in the middle which is assumed to propagate along the centerline of the cell at a constant speed $v_b$. (Bottom) Schematic picture of a spatial profile of preferred curvature $\kappa(s,t)$ (left) or preferred twist $\tau(s,t)$ (right) along the arclength $s$ of a filamentous cell.

Here, $\kappa_0(> 0)$ is the magnitude of the preferred curvature, $\xi$ is the interface thickness along $s$ connecting the regions of the different preferred curvatures, and $D$ defines the size of the activated section. As will be shown below, the experimentally observed patterns can be reproduced by appropriately choosing the parameters values of $\kappa_0$ and/or $D$.

For *Spiroplasma*, we assume that the preferred curvature is uniform, i.e., $\kappa$ is constant along the cell, while the preferred twist changes in space and time, $\tau_0(s,t) = \tau_0\psi(s - v_b t)$.

**Numerical scheme**

In the numerical simulation of our mathematical model sketched above, a rod is discretized into the chain of straight segments of length $a$, and a discrete orthogonal director frame, $\hat{d}_{\alpha,i}$, ($\alpha$ = 1,2,3), is assigned at each node $i$, where $i = 1,2,\cdots,N$. The bending and twisting forces derived from the appropriately discretized version of $E_{rod}$ have been previously described in detail, e.g., in Ref. [69], and will not be repeated here. In short, we employ the Euler angle representation to describe the configuration of the discrete director frames at each node. The Kirchhoff strain vector components in the discretized model, $\Omega_{a,i}$, and the corresponding discretized elastic energy $E_{\mathrm{rod}} = \sum_i E$ , with $E = E(\Omega_{1,i}, \Omega_{2,i}, \Omega_{3,i})$, can be given in terms of the three Euler angles. The variations of the elastic energy, $\delta E_{\mathrm{rod}}$, is related to the variations of the position of the node, $\mathbf{r}_i$, and (virtual) axial rotational angle about the local axis $\mathbf{d}_{3,i}$, $\phi_i$, through the variations of the corresponding Euler angles, from which we find the bending and twisting forces acting on the each node of the chain.

Note that, in addition to $E_{\mathrm{rod}}$, the elastic energy of the rod also includes the stretching energy penalty with a large stretching modulus that limits any bond extensibility within a few percent approximately ensuring the inextensibility condition implied in Eq. (1). In our present model, there are two more additional potentials, i.e., a short-ranged repulsive potential $E_{\mathrm{LJ}}$ that prevents the self-crossing of the rod-like cell and a short-range attractive potential between the rod centerline and the substrate that mimics a non-specific moderate adsorption of the cell surface onto the substrate in our experimental conditions.

**Loop formation dynamics**

Here we develop a simple scaling argument to discuss the dynamics of loop formation observed in *Haloplasma* cells. See also Fig. 4F. In our experimental measurements, the typical time to form a single loop is found to be $\tau_{\mathrm{loop}} \sim 1$ s. Suppose that a section of length $\Delta\ell \sim 2\pi R$, where $R$ is a radius of a loop formed in a rod-like cell of total length $L$, has a preferred curvature of magnitude $\kappa_0 \sim 1/R$, where $L \gg \Delta\ell$. Then, the straight configuration of the section of $\Delta\ell$ now has a deformation energy of $\delta U \sim A\kappa_0^2\,\Delta\ell \sim A\,\Delta\ell/R^2$, which drives the cell shape towards a new equilibrium (i.e., a looped) configuration. To make a loop of radius $R$, the section must pull additional lengths of $2\pi R$, requiring translational motions of the entire cell of length $L$ in a viscous medium. Let $\tau_{\mathrm{loop}}$ be the characteristic time to complete the loop formation, then the typical sliding velocity of the rod is $v \sim (\Delta l)/\tau_{\mathrm{loop}}$. The total power consumption, or the dissipated energy due to the frictional forces from the surrounding medium, is thus $Q \sim \gamma v^2 L \tau_{\mathrm{loop}} \sim \gamma(\Delta\ell)^2 L/\tau_{\mathrm{loop}}$, where $\gamma = 2\pi\eta$ is the friction coefficient of a slender object in a fluid of viscosity $\eta$. At steady state, we expect the energy balance $\delta U \sim Q$, leading to the scaling relation for $\tau_{\mathrm{loop}}$ given by

$$\tau_{\mathrm{loop}} \sim \frac{2\pi\gamma R^3 L}{A} \sim \frac{4\pi^2\eta R^3 L}{A}. \tag{3}$$

Assuming that the bending rigidity of *Haloplasma* should be similar to that of *Spiroplasma*, we use the recently reported values $A = 0.15 - 0.25$ pN$\cdot\,\mu$m$^2$ [68]. Plugging typical values characterizing cell size, $a \sim 200$ nm, $L \sim 10\ \mu$m, and using the water viscosity $\eta \sim 10^{-3}$ Pa$\cdot$ s, we obtain from Eq. (3) $\tau_{\mathrm{loop}} \sim 0.5$–$0.9$ s, in good agreement with our measurement. Therefore, the above argument overall validates the fundamental assumptions of our biophysical model, together with the parameter values used in the numerical simulations.

**Viscous dynamics and simulation parameters**

Due to the smallness of the system, the dynamics is overdamped, in which the frictional force from the surrounding viscous medium balances locally and instantaneously the sum of the internal forces (plus the force from the substrate) at each point of the rod. To mimic thermal agitations, we also include random forcing terms consistent with the fluctuation-dissipation relations in the background medium of temperature $T$. The resulting Langevin equations for $\mathbf{r}_i$ and $\phi_i$, where $i = 1,2,\cdots,N$, are rescaled appropriately (see below), and numerically integrated via the explicit Euler method with non-dimensional time steps typically $2.5 \times 10^{-5}$ that ensures a sufficient numerical accuracy. At each time, the director frames at each node are also updated, and the corresponding Euler angles are calculated for a new configuration of the rod centerline.

In the rescaling, the unit of length, energy and time are chosen, respectively, as the diameter of the cell, $a \approx 200$ nm, the thermal energy $k_BT \approx 4.0$ pN· nm, and the diffusion time $\tau = a^2/(\mu k_BT) \approx 1.8 \times 10^{-2}$ s, where $\mu = 1/(3\pi\eta a)$ is the self-mobility in the Stokes fluid. Experimentally, we have measured the typical velocity of the bending kink propagation in *Haloplasma* as $v_b = 5 - 15\mu$m/s (Fig. 4F). The corresponding rescaled velocity is thus set as $\hat{v}_b = v_b(\tau/a) = 0.5 - 1.5$. Consistent with the scaling argument above, we set the nondimensional bending modulus as $\hat{A} = A/(ak_BT) = 10^3$. We assume the twist-bend rigidity ratio $C/A = 0.667$, which corresponds to the Poisson's ratio 0.5 as valid to incompressible materials. The stretching modulus is chosen as $K = 16A/a^2$, which is valid to for an isotropic elastic rod and is sufficiently large to make the bond length variations negligibly small. The number of the nodes in the simulations is $N = L/a = 50 - 241$. The values of the relevant parameters for different types of observed motilities are summarized in table S3.

**Parameter values**

Table S3: The important non-dimensionalized parameter values used in our numerical simulations of the different types of the motilities.

| | $N$ | $\hat{v}_b$ | $\hat{\kappa}_0$ | $\hat{\tau}_0$ | $\hat{\xi}$ | $\hat{D}$ |
|---|---|---|---|---|---|---|
| *Haloplasma* kink | 241 | 1.38 | 0.5 | 0.1 | 0.4 | 5.65 |
| *Haloplasma* loop | 241 | 1.38 | 0.5 | 0.1 | 0.4 | 11.3 |
| *Spiroplasma* kink | 81 | 0.4 | 0.27 | 0.39 | 0.4 | 40.0 |

**References**


(18) H. Wada and R. R. Netz, Phys. Rev. Lett. **99**, 108102 (2007).
(65) R. E. Goldstein, A .Goriely, G. Huber and C. W. Wolgemuth, Phys. Rev. Lett. **84**, 1631 (2000).
(66) J. Yang, C. W. Wolgemuth, and G. Huber, Phys. Rev. Lett. **102**, 218102 (2009).
(67) C. Esparza L´opez and E. Lauga, Phys. Rev. Fluids **5**, 093102 (2020).
(68) G. Chirico and J. Langowski, Biopolymers **34**, 415 (1994); G. Chirico, Biopolymers **38**, 801 (1996).
(69) P. M. Ryan, J. W. Shaevitz and C. W. Wolgemuth, Phys. Rev. Lett. **131**, 178401 (2023).

**Movie S1.**
Field image video of syn3B original and syn3B−Halo7 for 10 s as real-time.

**Movie S2.**
Nucleoids in syn3B−Halo7 visualized by Hoechst for 2.5 s.

**Movie S3.**
Movement types I to IV of syn3B−Halo7 and type V of *H*MreB B1–A1 presented in 10-times slower speed than the realtime through eight frames.

**Movie S4.**
Coiling movements of *Haloplasma* and syn3B−Halo7 cells presented for 0.5 s.

**Movie S5.**
syn3B cell movements by two or more *H*MreBs presented for 10 s, as 1st and 2nd parts of movie. *H*MreB C2–A3–B4 did not move.

**Movie S6.**
Non-motile syn3B cells with expressing *H*MreBs (1st part), and syn3B cells with expressing single *H*MreBs (2nd part), presented for 5 s.

**Movie S7.**
syn3B cell movements by *H*MreB A2 presented for 10 s.

**Movie S8.**
Cryo-electron tomography with rendering.

**Movie S9.**
Motility modeling of movements by *Haloplasma* and *Spiroplasma* MreBs

**Data set S1. (separate file)**
List of genomes, *mreB*s and DNA constructs used in this study.
